# Disruption of the Wnt-antagonist *Apc* in the pituitary stem cells drives the development of adamantinomatous craniopharyngioma

**DOI:** 10.1101/2025.09.28.679021

**Authors:** James Blackburn, Laura Gomez-Corral, James Nicholson, Ashutosh Rai, Jingyi Xue, Rachael Tan, Kalyana C. Gangaraju, Angelica Gualtieri, Charlotte Hall, Pinaki Dutta, Márta Korbonits, Federico Roncaroli, Shannon W Davis, Carles Gaston-Massuet

**Author notes:** Corresponding author Carles Gaston-Massuet. These authors contributed equally to this work (Laura Gomez-Corral, James Nicholson).

## Abstract

Adamantinomatous craniopharyngiomas (aCPs) are complex intracranial neoplasms that generally arise in the sellar and suprasellar region of the brain which affect the endocrine and nervous systems causing severe sequelae. Activating mutations resulting in degradation-resistant forms of β-catenin (*CTNNB1*) have been shown to be the main driver for many of these neoplasms. However, the underlying genetic driver for a proportion of these tumours is still unknown. Using murine transgenic models, we show that genetic disruption of the Wnt-antagonist and tumour suppressor, adenomatous polyposis coli (*Apc*), within the pituitary progenitors/stem cells, leads to pituitary tumours that closely resemble human aCPs. These tumours present classic histopathological hallmarks of aCPs, such as large deposits of wet keratin, stellar-reticular-like cells, large cystic components and clusters of accumulating nucleo-cytoplasmic β-catenin that are slow-dividing and exhibit a secretory phenotype. We show that a hypomorphic allele of *Apc* is sufficient for tumour development, indicating that disruption of *Apc* function can lead to aCP formation. Moreover, we identify that bi-allelic loss of *Apc* in the Sox2+ve pituitary stem cells is sufficient to initiate tumour formation, indicating that Sox2+ve stem cells are the cell origin of these *Apc*-driven aCPs. Transcriptomic analyses of early tumour-initiating cells revealed that *Apc-*driven clusters of accumulating β-catenin undergo senescence-associated secretory phenotype (SASP), which is p21-mediated and results in secretion of inflammasome, angiosome and developmental growth factors. Our data unequivocally show a causal role for the disruption of the tumour suppressor *Apc* as a primary driver of aCPs independent of mutations in β-catenin. We provide murine models representing a novel genetic subtype of human aCPs offering insights into aCP pathogenesis. Our work reinforces the importance of genetic testing for mutations in *APC* in patients with aCPs and identifies a potential need to screen patients with familial adenomatous polyposis (FAP) or spectrum of *APC*-pathogenic syndromes for aCPs in early life.

## Introduction

Craniopharyngiomas (CPs) are complex neoplasms classified by the World Health Organisation (WHO) as grade 1 benign epithelial tumours typically located within the sellar or suprasellar region of the brain^1^. They are classified into two distinct subtypes: papillary craniopharyngioma (pCPs) which account for around 10% and adamantinomatous craniopharyngioma (aCPs) representing the remaining 90% of all CPs^2^. Molecularly, 90% of pCPs are driven by mutations in BRAF p.V600E^3,4^, whilst the majority of aCPs are driven by mutations in the *CTNNB1* gene encoding for β-catenin. Although rare, with an incidence of 1.7 per 1,000,000^2,5^, aCPs are the most common suprasellar and non-neuroepithelial intracranial tumour in childhood and have been shown to originate from progenitors/stem cells of the pituitary gland^6,7^. aCPs can occur at any age^8^ but have a bimodal age of presentation between 5-14 and 55-70 years old^9^. Although classified as benign, aCPs often exhibit infiltrative behaviour, invading critical neighbouring structures such as the hypothalamus and optic and cranial nerves. This makes total surgical resection technically difficult resulting in a high rate of recurrence^2,9^. Current treatment of aCPs is limited to surgery combined with radiotherapy, both of which are associated with high morbidity and significant disabling side effects.

The prevalence of mutations in β-catenin in human aCPs varies from 30% to 100% depending on the study^10-16^. The variability in mutation frequency may be due to small patient cohorts and/or the use of sequencing methods lacking the high depth of reading required for the detection of low allelic frequency variants in these highly heterogeneous tumours. Consequently, main genetic drives for a proportion of aCPs remains unknown. Previous transgenic murine models have been instrumental to understanding aCP pathogenesis^6,7,17,18^. Pituitary tissue-specific expression of a degradation-resistant form of β-catenin in murine pituitary stem cells has demonstrated that mutations in β-catenin causally drive development of aCPs^6^. These tumours have similar histopathological and molecular features to human aCPs, including the presence of whorl-like clusters or clusters of accumulating nucleo-cytoplasmic β-catenin that overactivate the canonical Wnt/β-catenin pathway. These clusters have been shown both in mice^18^ and human^19,20^ to undergo oncogene-mediated senescence and act as secretory hubs to modify the tumour microenvironment^18,19^. β-catenin is the central component of the Wnt/β-catenin pathway, which is crucial for development and stem cell homeostasis, with its dysregulation leading to tumour formation^21-23^. In the absence of secreted Wnt ligands, β-catenin is phosphorylated by a destruction complex formed by the scaffold protein Axin2, adenomatous polyposis coli (APC) and the kinase, GSK-3β. At this stage, β-catenin is phosphorylated by GSK-3β, ubiquitinated and targeted for proteasomal degradation, thereby regulating the levels of active β-catenin present in the cell^24^. Mutations in negative regulators of the Wnt pathway, that function to degrade β-catenin, such as adenomatous polyposis coli (*APC*), result in increased Wnt/β-catenin signalling, and are involved in the pathogeneisis of numerous cancers^25^. Indeed, both somatic and germline mutations in *APC* are the main drivers of colon carcinoma. Germline mutations cause familial adenomatous polyposis (FAP)^26^ and the related spectrum of *APC*-pathogenic syndromes such as Gardner^27^ and Turcot Syndrome^28^ which are characterised by colonic adenomas alongside tumours within the brain, bone and skin^29,30^. Recently, an association between FAP and aCPs has been reported in 2 separate studies^31,32^. These reports, using whole exome sequencing, identified 3 cases of aCPs harbouring germline mutations in *APC*, implicating this negative regulator of Wnt/β-catenin pathway in the pathogenesis of aCPs. A further possible association between FAP and aCPs has been hypothesised as some patients with Gardner syndrome, which carry germline mutations in *APC*, have been reported to develop ectopic aCPs^33-39^. However, a causal role of *APC* mutations in the formation of these aCPs, could not be determined due to the lack of tumour genomic sequencing data for these patients^39-42^. Moreover, a recent study that analysed two transgenic models to genetically delete *Apc* concluded that genetic deletion of *Apc* in rodent pituitary progenitors was not tumorigenic resulting in the absence of aCP formation even in older mice^18^. Therefore, although an association between aCPs and mutations in *APC* exists, the causal role for the dysfunction of *Apc* in aCP pathogenesis has remained elusive so far.

To demonstrate the impact of *Apc* disruption in aCP pathogenesis, we used murine transgenic models combined with transcriptomic analyses of early aCP tumour-initiating cells to demonstrate a causal role of *Apc* disruption in aCP pathogenesis. We show that biallelic loss of the truncated form of *Apc* within pituitary progenitors, by deleting *Apc exon 15*, results in severe intrauterine aCPs that are not compatible with life. Generation of an *Apc* hypomorphic allele results in aCPs postnatally that accurately recapitulate aCP disease progression, presenting with key histological and molecular hallmarks of human aCPs, as well as clinically relevant tumour-induced hypothalamic obesity. Transcriptomic analyses of early tumour-initiating cells revealed that the disruption of *Apc* within pituitary stem cells leads to clusters of accumulating nucleo-cytoplasmic β-catenin that undergo a senescence associated secretory phenotype (SASP). We prove that this secretory signature consists of specific pro-inflammatory mediators, chemokines, cytokines and developmental growth factors. Importantly, these *Apc-* driven tumour-initiating cells exhibit transcriptional signatures that could be pharmacologically targeted. Our work demonstrates a direct causal effect of *Apc* disruption as a main driver of aCPs independent of β-catenin mutations. We provide further understanding of the underlying genetics of aCPs and present murine models representing a novel genetic subtype of human aCPs. Our results reinforce the need for genetic testing for mutations in *APC* in patients with aCP and the importance of increased surveillance for patients with FAP or *APC*-related syndromes for the development of extracolonic tumours, such as aCPs.

## Results

### Bi-allelic loss of *Apc exon 15* in the pituitary embryonic precursors leads to severe aCPs during embryogenesis

To determine the causal role of *APC* in craniopharyngioma formation, we employed a murine transgenic approach to delete exon 15 of *Apc* specifically in embryonic pituitary progenitor cells. We used the conditional *Apc* allele in which exon 15 has been flanked with *LoxP* sites *Apc*^*LoxP(Exon15)/LoxP(Exon15)* 43^ to generate a conditional truncated form of *Apc*. Exon 15 of *Apc* encompasses 2200 amino-acids, including the functional domain responsible for β-catenin degradation and the C-terminal domain involved in microtubule binding^43^. To conditionally delete *Apc* within embryonic pituitary progenitor cells, we used the *Prop1:Cre* driver^44,45^, that expresses Cre within pituitary progenitors from embryonic day (e) 10 of gestation, to generate homozygotes *Prop1:Cre;Apc* ^*LoxP(Exon15)/LoxP(Exon15)*^ (hereafter referred to as *Cre:Apc*^*LoxP15/15*^ for simplicity). Homozygotes *Cre:Apc*^*LoxP15/15*^ pups died perinatally, with none of the mutant pups reaching postnatal day 7. Dissections of pituitary glands revealed large tumours in all *Cre:Apc*^*LoxP15/15*^ embryos compared to their wild type (Wt, *Apc*^*+/+*^) littermates (**Fig. 1a**). Histological analyses using haematoxylin and eosin staining (H&E) and antibodies against terminally differentiated hormone producing cells, revealed that *Cre:Apc*^*LoxP15/15*^ mutant embryos exhibit large pituitary tumours that are almost devoid of all terminally differentiated hormone producing cells (**Fig. 1b**). These tumours are characterised by aCP-histopathological features such as wet keratin and whorl-like structures consisting of cell clusters of nucleo-cytoplasmic positive β-catenin (**Fig. 1d**). Abnormal larger pituitary glands were observed as early as embryonic day (e) 11 and progressively worsen throughout gestation, exhibiting statistically significant increase in pituitary volume (**Fig. 1c**) secondary to increased mitotic index (MI) (**Fig. 1 e-f**). To further characterise these tumours, we performed double immunofluorescence with β-catenin and the proliferation marker phospho-histone H3 (α-pHH3) or the pituitary stem cell markers Sox2 or Sox9, as aCP-histological hallmarks^6,17,46,47^. Double immunofluorescence at e13.5 and e18.5 revealed large clusters of nucleo-cytoplasmic accumulating β-catenin, which were negative for pHH3. Notably most of the dividing cells were located surrounding the β-catenin+ve cell clusters (**Fig. 2 a-b**). As previously described in β-catenin-driven aCP tumours^6,7^, these β-catenin+ve clusters express the pan-pituitary stem cell marker Sox2 (**Fig. 2 c-d**), but are negative for the pituitary transcription factor Sox9. Instead, Sox9+ve cells were localised around the β-catenin+ve cell clusters (**Fig. 2 e-f**). We confirmed the efficiency of recombination of the exon 15 of *Apc* by RT-qPCR and identified 100% deletion of exon 15 by our *Prop1:Cre* driver line (**Supplementary Fig. 1d**). Together, our results demonstrate that bi-allelic loss of *Apc* exon 15 within pituitary progenitor cells, results in intrauterine aCPs associated with early postnatal lethality due to tumour size and severity, with absence of differentiation of hormone secreting cells in the anterior gland.

**Fig 1.**
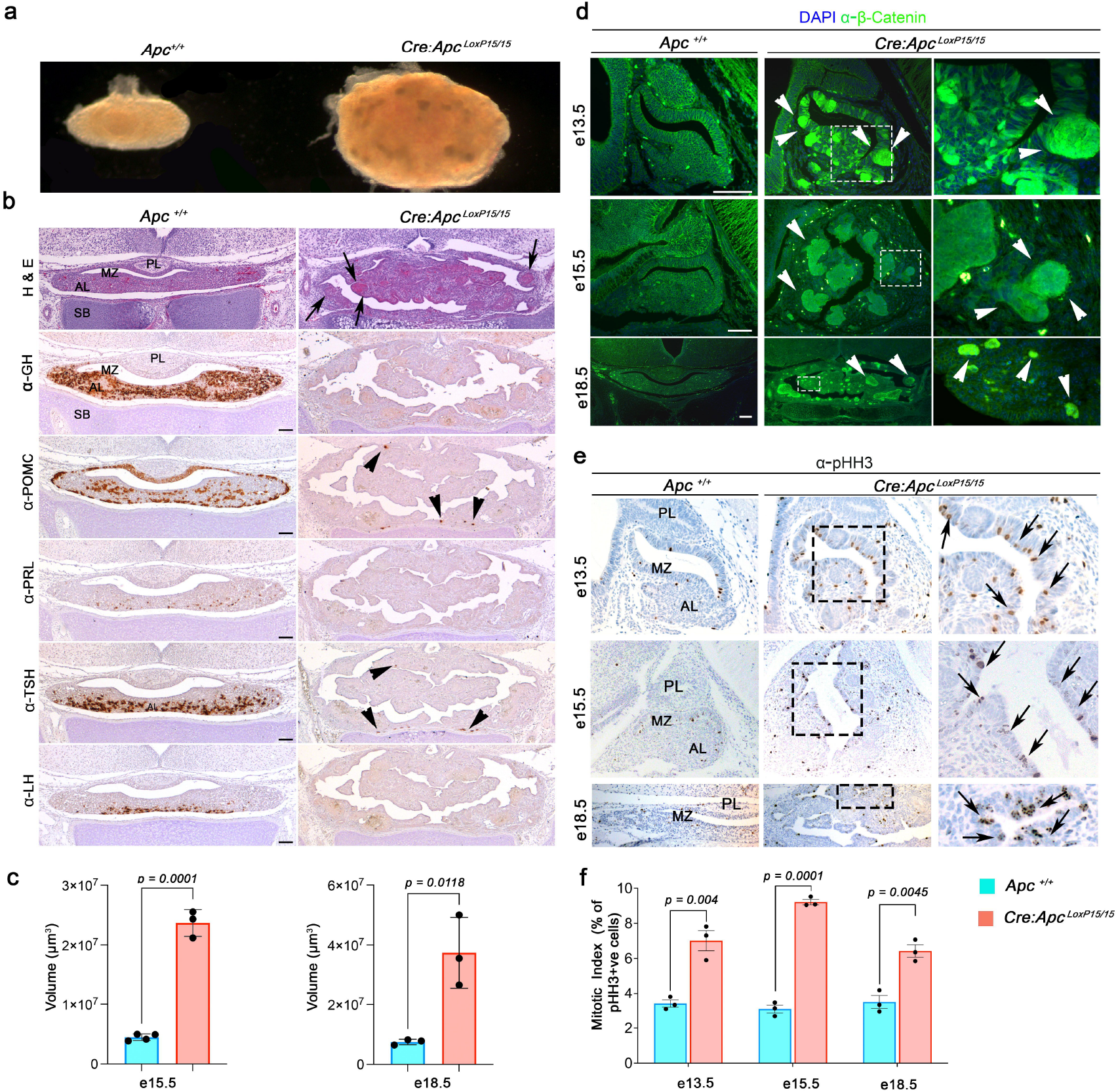
Genetic deletion of *Apc exon15* in the pituitary progenitors results in large embryonic non-secreting pituitary tumours with classic histological hallmarks of aCPs. **a**) Wholemount picture of dissected pituitaries at embryonic day e18.5 of a wild type (Wt, *Apc*^*+/+*^) compared to mutant *Cre;Apc*^*LoxP15/15*^ exhibiting a large tumour. **b**) Coronal sections through the pituitary gland at e18.5 stained with H&E reveal a large hyperplastic tumour with epithelial whorl-like structures (arrows) within the tumour parenchyma in the *Cre;Apc*^*LoxP15/15*^ (right column) compared to Wt pituitary glands (left column). IHC against GH, POMC (melanotrophs and ACTH), PRL, TSH and LH reveals that the *Cre;Apc*^*LoxP15/15*^ tumours are largely devoid of hormone-producing cells, apart from some scattered positive foci of POMC and TSH (arrowheads). **c**) Volumetric quantifications indicate that the *Cre;Apc*^*LoxP15/15*^ mutant pituitaries have statistically significant larger volumes compared to Wt littermates at both e15.5 and e18.5 of gestation. **d**) Immunofluorescence against β-catenin reveals clusters of nucleo-cytoplasmic β-catenin accumulating cells which are hallmarks of aCPs in the developing pituitary gland at e13.5, e15.5 and e18.5 in the mutant *Cre;Apc*^*LoxP15/15*^ (white arrowheads) compared to the membranous only β-catenin staining in the Wt pituitaries. **e**) IHC against the proliferation maker, phospho-histone H3 (α-pHH3), reveals increase in pHH3-postive cells in the *Cre;Apc*^*LoxP15/15*^ mutant pituitaries at e13.5, e15.5 and e18.5 compared to Wt. Quantification of the percentage of pHH3+ve cells revealed a statistically significant increased mitotic index (MI) in *Cre;Apc*^*LoxP15/15*^ compared to their Wt littermates. Right-hand column images in **d & e** are enlarged images of the squared boxes of the images in the central column. *P* values in **c** and **f** were obtained using unpaired two-tailed Student’s *T*-test and data represented as mean ± SEM, from 3/4 pituitaries per genotype. Images are representative of 4-5 embryos per genotype. Abbreviations: AL, anterior lobe; LH, luteinizing hormone; GH, growth hormone; H&E, haematoxylin and eosin; IHC, immuno-histochemistry; MZ, marginal zone; PL, posterior lobe; POMC, Pro-opiomelanocortin; PRL, prolactin; SB, sphenoid bone; TSH, thyroid-stimulating hormone. Scale bar in **b** represents 50 μm; and in **d** represenst 200 μm.

**Fig 2.**
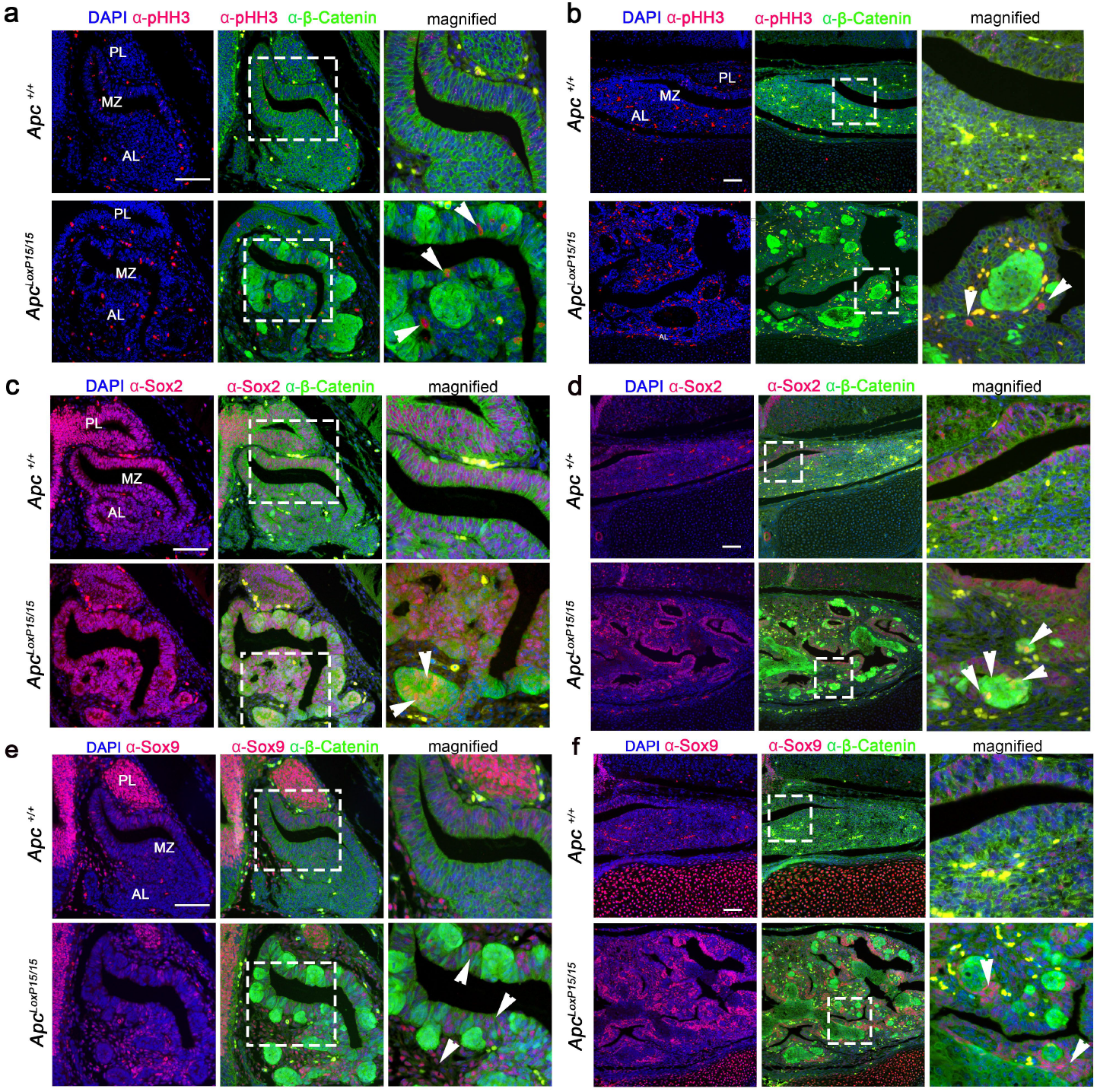
Genetic deletion of *Apc exon15* (*Cre;Apc*^*LoxP15/15*^) leads to aCP histological hallmarks such as cell clusters of accumulating nucleo-cytoplasmic β-catenin. These express the stem cell marker Sox2, are negative for the Sox9 and have low proliferative capacity. **a-f**) Histological sections through the pituitary gland at e13.5 (**a, c, e**) and e18.5 (**b, d, f**) immune-stained with α-pHH3 (**a, b**), α-Sox2 (**c, d**) and α-Sox9 (**e, f**). **a, b**) Double immunofluorescence against pHH3 (red) and β-catenin (green) reveals the presence of clusters of accumulating nucleo-cytoplasmic β-catenin (green) which do not express the proliferation marker pHH3 in the *Cre;Apc*^*LoxP15/15*^. Proliferating cells (pHH3+ve cells, white arrowheads) were always found outside of the β-catenin cell clusters at e13.5 (**a**) and e18.5 (**b**). **c, d**) Double immunofluorescence against β-catenin (green) and α-Sox2 (red) reveals that clusters of accumulating β-catenin express the pituitary stem cell marker Sox2 (white arrowheads). **e, f**) Double immunofluorescence against β-catenin (green) and α-Sox9 (red) shows that clusters of accumulating β-catenin do not express transcription factor Sox9 but rather, the Sox9+ve cells are found surrounding the β-catenin+ve clusters. Note that no clusters of β-catenin were observed in any of the Wt pituitaries (*Apc*^*+/+*^) with β-catenin being localised to the cell membrane. Magnified images in the right columns in **a** to **f** represent the white-dashed squared areas in the middle column panels. Images are representative of 4 embryos per genotype per each gestational stage. Abbreviations: AL, anterior lobe; MZ, marginal zone; PL, posterior lobe. Scale bars in **a, c, e** represent 200 μm and in **b, d, f** represent 100 μm.

### A hypomorphic allele for *Apc* results in postnatal aCP tumour formation

To overcome the early lethality observed in the *Cre:Apc*^*LoxP15/15*^ animals, we took a genetic approach to generate another mutant allele of *Apc*. To achieve this, we utilised the *Apx*^*LoxP(Exon1-15)*^ line in which *LoxP*-sites have been introduced to flank all the *Apc* exons, from exon 1 to exon 15, enabling deletion of the entire *Apc* locus^48^. Homozygotes *Prop1:Cre*:*Apc*^*LoxPExon 1-15/LoxPExon1-15*^ (hereby referred to as *Cre:Apc*^*LoxP1-15/1-15*^ for simplicity) exhibited enlarged pituitary glands due to increased MI (**Supplementary Fig. 1 a-b**) with decreased number of growth hormone-producing cells (**Supplementary Fig. 1 a**). However, no tumour formation occurred in any of the mutant mice (*Cre:Apc*^*LoxP1-15/1-15*^) analysed after 2 years, other than hyperplasia of the anterior pituitary gland (data not shown). This prompted us to investigate the efficiency of Cre recombination of exons *1-15* at the *Apc* locus by RT-qPCR in our homozygote *Cre:Apc*^*LoxP1-15/1-15*^ pituitaries. Importantly, we identified that 16.37% of Wt *Apc* exon 15 mRNA was being produced, indicating inefficient recombination of the *Apc* locus (**Supplementary Fig. 1d**). This result indicates that a small amount of Wt *Apc* mRNA is sufficient to prevent tumour formation, resulting in pituitary hyperplasia instead of tumours. Hence, we reasoned that further reduction of Wt *Apc* levels by combining the *Cre:Apc*^*LoxP15/+*^ (with 100% recombination efficiency) with the *Cre:Apc*^*LoxP 1-15/+*^ alleles to generate a compound heterozygote *Cre*;*Apc*^*LoxP1-15/15*^ could result in a milder phenotype compared to the *Cre:Apc*^*LoxP15/15*^. Indeed, all *Cre:Apc*^*LoxP1-15/15*^ compound mutants survived to adult compared to the early perinatal lethality observed in the *Cre:Apc*^*LoxP15/15*^ pups (**Fig. 1**). All *Cre:Apc*^*LoxP1-15/15*^ compound mutants developed large pituitary tumours postnatally, leading to tumour-associated co-morbidities such as severe obesity and reduced life expectancy (**Fig. 3 a-f**). Obesity secondary to tumour formation was apparent in all mutant *Cre:Apc*^*LoxP1-15/15*^ mice from 4 months of age(**Fig. 3 c-d**), an important phenotype that has not being observed in previously published murine aCP models^6^. Histological analyses of these tumours revealed the presence of large cystic components, stellar reticular-like cells and nucleo-cytoplasmic β-catenin+ve cell clusters, all of which are histological diagnostic hallmarks of aCPs (**Fig. 3b**). Interestingly, Kaplan Meier survival curves revealed a gender difference in survival, with female mice surviving longer than males (**Fig. 3 e-f**).

**Fig 3.**
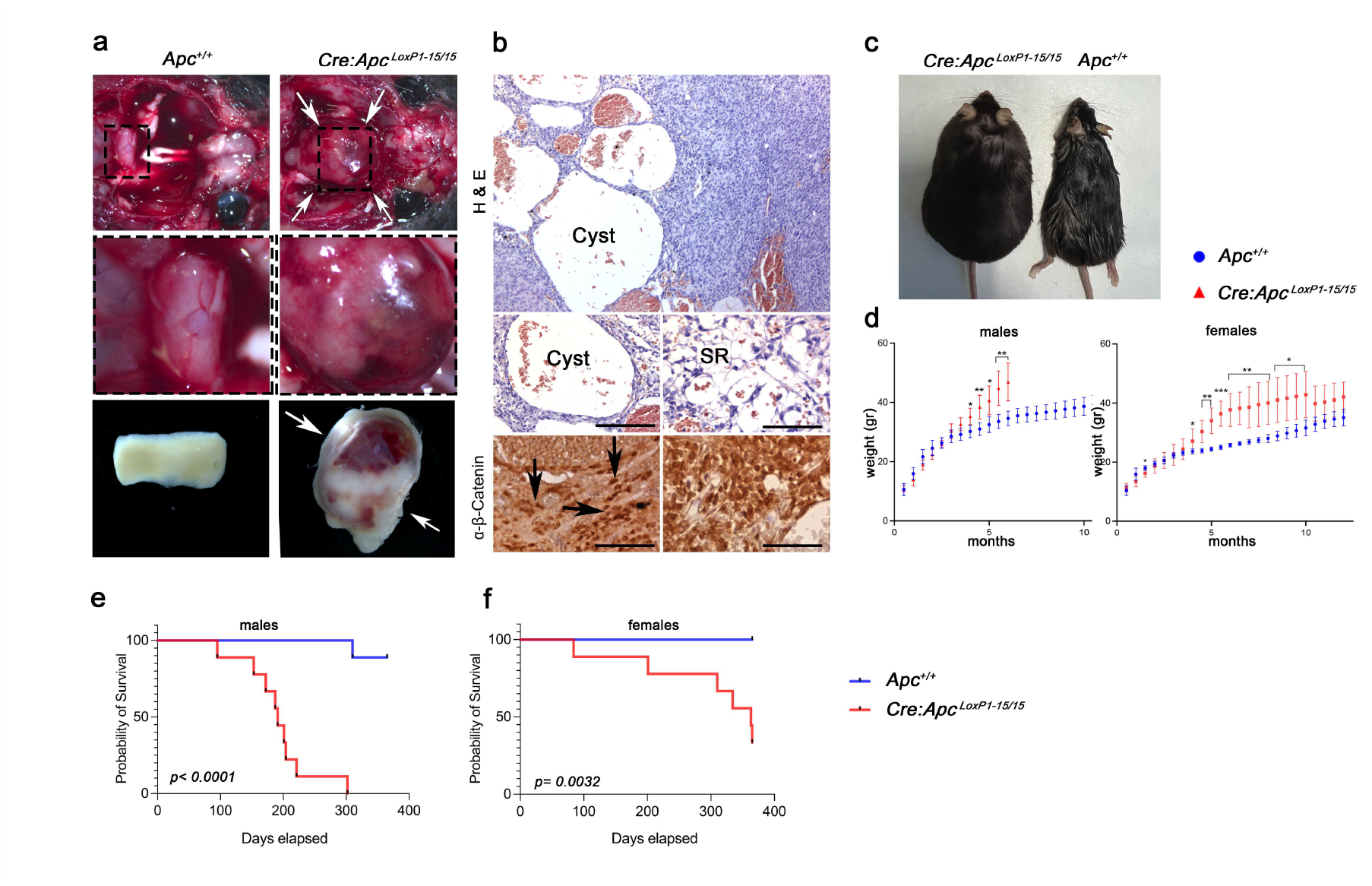
A hypomorphic allele of *Apc* (*Cre:Apc*^*LoxP1-15/15*^) develops large pituitary tumours postnatally. **a**) Picture after removal of the scalp and brain of 6 month old *Cre:Apc*^*LoxP1-15/15*^ mutants revealed pituitary tumours (white arrows). **b**) H&E sections of the tumours reveal large cystic components (Cyst) and some cells reminiscent of stellate reticulum (SR) cells. IHC against β-catenin shows cells with nuclear positivity within some cell clusters (black arrows). **c-d**) Male and female hypomorphic animals exhibit statistically significant weight gain starting from 4 months of age, with all mutant animals exhibiting obesity compared to their Wt littermates (**d**). Kaplan Meier survival curves from males (**e**) and females (**f**) show that hypomorphic *Cre:Apc*^*LoxP1-15/15*^ mutant animals have shorter lifespans compared to their Wt littermates. Note that males (**e**) have a shorter life span compared to females (**f**), with a 50% probability of survival at 6 months (180 days), whilst females exhibited a longer life span (320 days). *P* values in **d** represent ***<0.001; ** <0.01 and were obtained using unpaired two-tailed Student’s *T*-test and data represented as mean ± SEM. **e & f** *P* values of survival curves were obtained by a long-rank Mantel-Cox test. Abbreviations: H&E, haematoxylin and eosin; SR, stellate reticulum cells. Scale bars in **b** represent 100 μm.

Characterisation of tumours at 3 months of age also revealed all the typical histological diagnostic hallmarks of human aCPs including cysts, nucleo-cytoplasmic β-catenin+ve cell clusters, large foci of wet keratin and increased MI (**Fig. 4 a-e**). Immunostaining of the pituitary gland for terminally differentiated hormones, revealed the absence of hormone-producing cells within the developing tumour in keeping with the non-hormone secreting nature of aCPs (**Fig. 4d**). H&E staining revealed wet keratin deposits, cysts and nucleo-cytoplasmic β-catenin+ve cell clusters often surrounding the wet keratin foci (**Fig. 4e**). To analyse the early tumour characteristics of the *Cre:Apc*^*LoxP1-15/*15^ hypomorphs at e18.5, double immunofluorescence against β-catenin and pHH3, Sox2 or Sox9 was performed. These early tumours exhibit clusters of accumulating nucleo-cytoplasmic β-catenin, which are negative for pHH3, with dividing cells localised to the surrounding of β-catenin+ve cell clusters (**Fig. 5a**). In contrast with the perinatal lethal *Cre:Apc*^*LoxP15/15*^ model, the β-catenin+ve clusters of the *Cre:Apc*^*LoxP1-15/15*^ tumours were smaller (**Supplementary Fig. 2**). The β-catenin+ve cell clusters were all positive for the pituitary stem cell marker Sox2, and negative for Sox9, with Sox9+ve cells localised around the β-catenin+ve cell clusters (**Fig. 5a**). Clonogenic assays found a statistically significant increase in the number of stem cell colonies and the number of cells per colony in mutant *Cre:Apc*^*LoxP1-15/15*^ derived pituitary stem cells at e18.5 compared to Wt pituitaries, indicating that mutant derived pituitary stem cells have more proliferative capacity (**Fig. 5b**). At this embryonic stage, IHC for terminally differentiated hormones revealed a decrease in hormone-producing cells compared to Wt with a clear decrease in GH, TSH, FSH and LH, whilst no differences were observed in other hormones (**Fig. 5c**). Next, RT-qPCR was used to identify the level of recombination in the *Cre:Apc*^*LoxP1-15/15*^ compound mutants. This identified that Wt *Apc exon15* mRNA was present in small amounts (7.5% compared to Wt cells) whilst the exon 15 from the *Cre:Apc*^*LoxP15/15*^ was fully deleted (**Supplementary Fig. 1d**). This result indicates that the *Cre:Apc*^*loxP1-15/15*^ expresses small amounts of Wt Apc and is therefore the hypomorphic allele of Apc. These low levels of Wt *Apc* are insufficient to prevent tumour formation in compound *Cre:Apc*^*LoxP1-15/15*^ animals, but result in a milder phenotype compared to *Cre:Apc*^*15/15*^ embryonic aCPs which exhibits 100% recombination of the *Apc-exon15* and are not viable postnatally.

**Fig 4.**
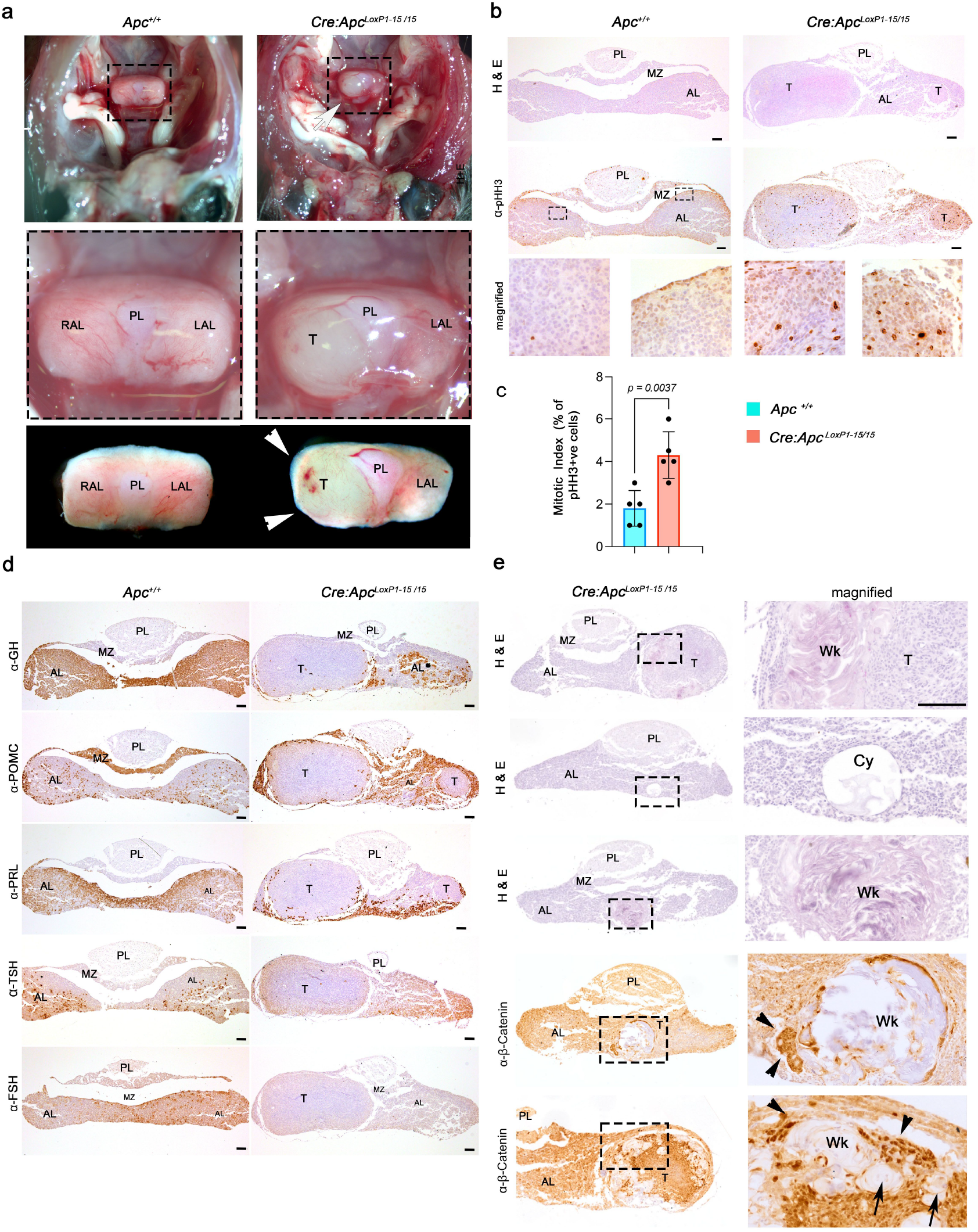
Mid-stage tumours from *Cre:Apc*^*LoxP1-15/15*^ hypomorphs exhibit classic histological hallmarks of aCPs. **a**) Picture after removal of the scalp and brain in 3 month old, *Cre:Apc*^*LoxP1-15/15*^ mutants revealed hyperplastic pituitaries with small tumours (white arrows & T). **b**) H&E staining of coronal sections through the *Cre:Apc*^*LoxP1- 15/15*^ mutant pituitaries revealed developing tumours within the pituitary parenchyma of differing sizes (marked as T). These tumours have increased pHH3+ve cells, resulting in a higher mitotic index (MI) (**c**). **d**) IHC against GH, POMC (melanotrophs and ACTH), PRL, TSH and FSH reveals that the developing tumours are devoid of hormone-producing cells. **e**) H&E staining of coronal sections through *Cre:Apc*^*LoxP1-15/15*^ mutant pituitaries revealed histological hallmarks of aCPs, such as deposits of wet keratin (Wk) and small cysts (Cy). IHC against β-catenin in mutant pituitaries revealed the presence of cell clusters of nucleo-cytoplasmic β-catenin (black arrowheads) surrounding nodules of wet keratin (Wk, black arrows). *P* value in (**c**) was obtained using unpaired two-tailed Student’s *T*-test and data represented as mean ± SEM. Images are representative of 5 mutant pituitaries per genotype. Abbreviations: AL, anterior lobe; FSH, follicle stimulating hormone; GH, growth hormone; PL, posterior lobe; PRL, prolactin; POMC, pro-opiomelanocortin; MZ, marginal zone; RAL, right anterior lobe; LAL, left anterior lobe; T, tumours; TSH, thyroid-stimulating hormone; Wk, wet keratin. Scale bars in **b & d** represent 100 μm, and in **e** represents 50 μm.

**Fig 5.**
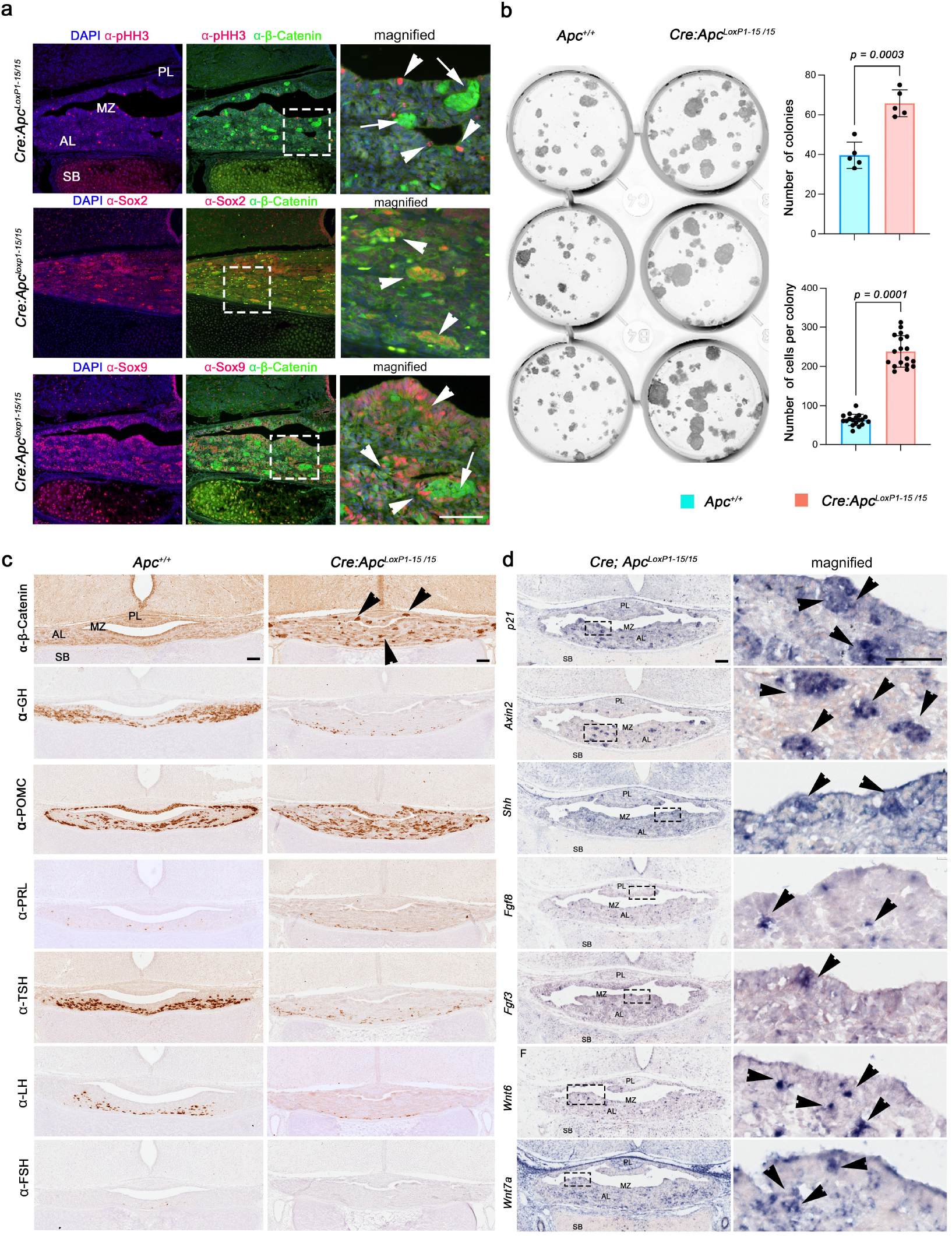
Embryonic pituitaries from *Cre:Apc*^*LoxP 1-15/15*^ embryos exhibit all the features of early aCP development. **a**) Double immunofluorescence of coronal sections through the developing pituitary gland from the *Cre:Apc*^*LoxP 1-15/15*^ embryos at e18.5 against pHH3, Sox2 and Sox9 with β-catenin. Clusters of nucleo-cytoplasmic β-catenin +ve cells are present throughout the pituitary parenchyma (green, white arrows) with pHH3+ve cells (red, white arrowheads) scattered around the clusters of β-catenin (white arrows in the magnified image). All clusters of accumulating β-catenin express the pituitary stem cell marker Sox2 (red, white arrowheads) and the Sox9+ve cells are located around the β-catenin+ve clusters. The right column images are magnified images from the white squared areas in the middle column. **b**) Pituitary stem cell culture from hypomorphic *Cre:Apc*^*LoxP 1-15/15*^ mutants exhibit higher clonogenic potential evidenced by both increased number of colonies and number of cells per colony compared to Wt pituitary stem cells. **c**) IHC against β-catenin in coronal sections through the pituitary and brain at e18.5 revealed clusters of cells of accumulating β-catenin in *Cre:Apc*^*LoxP 1- 15/15*^ (right column, black arrowheads) compared to Wt (left column). IHC against GH, TSH, LH and FSH reveals that *Cre:Apc*^*LoxP 1-15/15*^ hypomorphic mutant pituitaries exhibit a reduction in these hormone-producing cells compared to their Wt littermates, whilst POMC (melanotrophs and ACTH) and PRL show similar expression between genotypes. **d**) *In situ* hybridisation on coronal sections through the pituitary gland of *Cre:Apc*^*LoxP1-15/15*^ mutants reveals upregulation of the canonical downstream Wnt-effector *Axin2* and the senescence marker *p21* in a cell cluster fashion. The developmental factors *Shh, Fgf8, Fgf3* and *Wnt6, Wnt7a* were highly upregulated within cell clusters throughout the pituitary parenchyma. Right column panels are magnified images from the dotted squared areas in the left column. *P* values in **b** were obtained using unpaired two-tailed Student’s *T*-test and data represented as mean ± SEM. Images are representative of 5 embryos per genotype. Abbreviations: AL, anterior lobe; LH, luteinising hormone; GH, growth hormone; IHC, immuno-histochemistry; MZ, marginal zone; PL, posterior lobe; PRL, prolactin; POMC, Pro-opiomelanocortin; SB, sphenoid bone; TSH, thyroid-stimulating hormone. Scale bars in **b** represent 50 μm; and in **c** and **d** represent 100 μm.

β-catenin-mutated+ve aCPs are characterised by the presence of clusters of accumulating nucleo-cytoplasmic β-catenin that undergo senescence secretory associated phenotype (SASP) and secrete factors into the tumour microenvironment^18,19,49^. We, therefore, analysed by *in situ* hybridisation the expression of some markers of β-catenin+ve clusters previously reported in mouse and human aCPs^6,17^. All clusters analysed from the *Cre:Apc*^*loxP1-15/15*^ hypomorphic pituitaries expressed the canonical Wnt-activating target, *Axin2*, indicating β-catenin+ve cell clusters over-activate the Wnt canonical pathway. Moreover, these clusters over-express the developmentally secreted factors *Wnt6, Wnt7a, Fgf3, Fgf8* and *Shh* in a cluster fashion (**Fig. 5d**). Importantly, these clusters upregulate the senescence associated marker *Cdkn1* (*p21*) in accordance with activation of SASP described in both human^19,49^ and murine aCPs^17,18^. Taken together, our results demonstrate that a hypomorphic allele of *Apc* leads to tumour formation postnatally, exhibiting all the characteristic histological and molecular features of aCP tumours, including associated comorbidities such as tumour-induced obesity. These results further demonstrate a causal role of *Apc* dysfunction in the development of aCPs.

### Biallelic loss of *Apc exon 15* in the Sox2+ve pituitary stem cells is sufficient to initiate aCP tumour formation

Previous studies using a conditional transgenic line have shown that Sox2+ve pituitary stem cells bearing oncogenic β-catenin (*Ctnnb1*^*LoxpExon3/*+^) can initiate aCP tumour formation^7^. To further prove a causative role of *Apc* in aCP formation, we employed an inducible *Sox2*^*CreERT2/+*^ transgenic line^50^ in which tamoxifen (Tam) induced Cre is achieved within Sox2+ve stem cells. To enable the deletion of *exon 15* of *Apc* we crossed the *Sox2*^*CreERT2*/+^ to the *Apc*^*LoxP(Exon15)/LoxP(Exon15)*^ strain to generate *Sox2*^*CreERT2/+*^*;Apc*^*LoxP(Exon15)/LoxP(Exon15)*^ embryos (hereafter refer to as *Sox2*^*CreERT2/+*^*;Apc*^*LoxP15/15*^ for simplicity). Deletion of *Apc exon 15* was achieved with 3 consecutive daily injections of Tam at 0.1 mg/gr/mouse during pregnancy starting at e12.5, and embryos were analysed at e18.5 (**Fig. 6a**).

**Fig 6.**
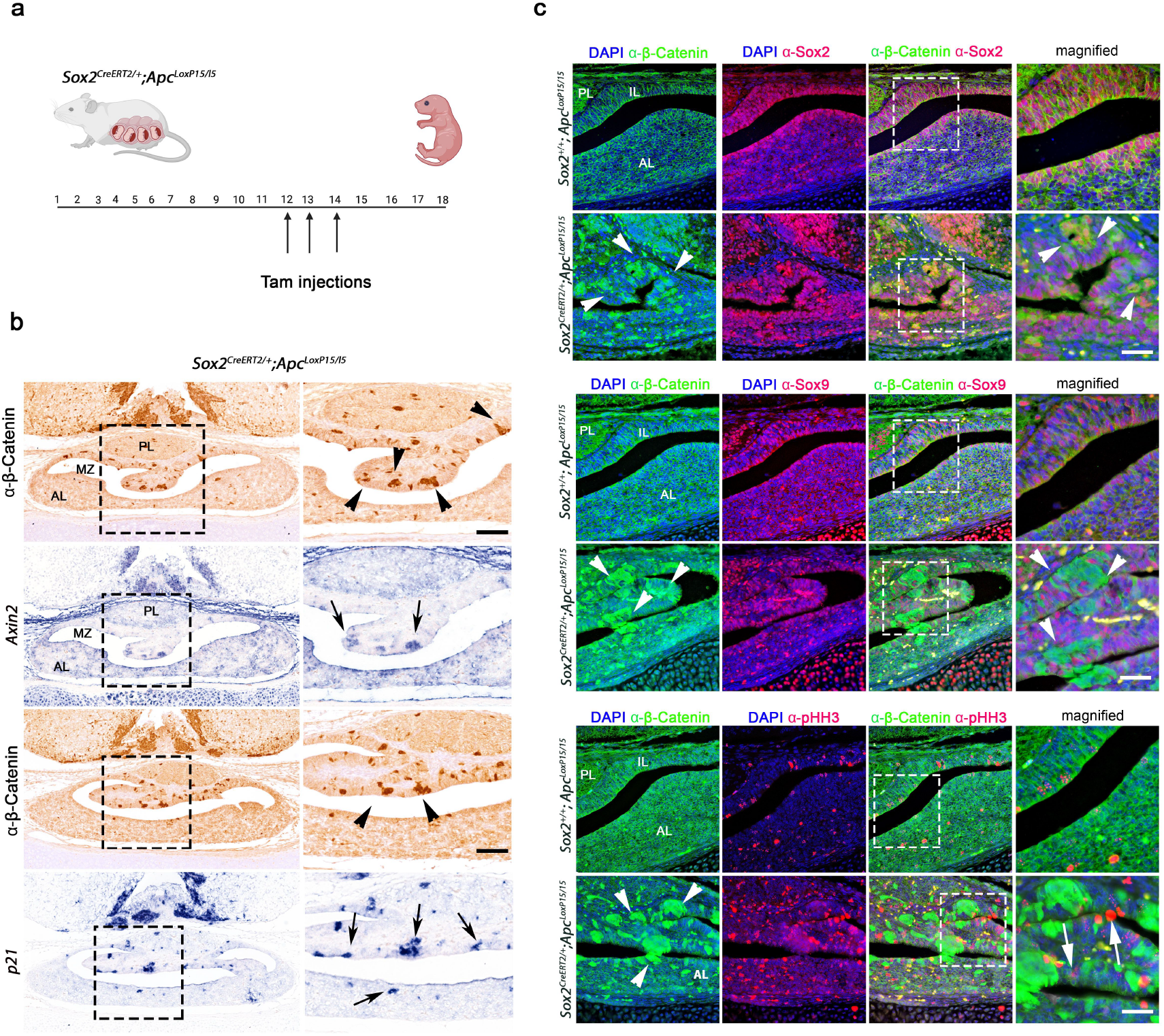
Deletion of *Apc exon15* in the Sox2+ve pituitary stem cells results in early aCP formation. **a**) Experimental diagram: to achieve deletion of *Apc exon15* (*Sox2*^*CreERT2/+*^*;Apc*^*LoxP15/15*^), pregnant females were Tam injected (0.1 mg/g of body weight) with 3 consecutive daily injections at days e12, e13 and e14 of pregnancy. Embryos were dissected at e18.5 and pituitary glands were analysed by *in situ* hybridisation and immunostaining. **b**) IHC against β-catenin revealed the presence of multiple cell clusters of nucleo-cytoplasmic accumulating β-catenin (arrowheads). *In situ* hybridisation on consecutive sections for *Axin2* and the senescence marker *p21* reveals the upregulation of *Axin2* and *p21* within the β-catenin positive clusters (arrows). Pictures in the right column are magnified images from the squared dashed boxes in the left column. **c**) Double immunofluorescence against Sox2, Sox9, pHH3 (red) and β-catenin (green) revealed that Tam-induced *Sox2*^*CreERT2/+*^*;Apc*^*LoxP15/15*^ embryos develop early aCP lesions characterised by clusters of nucleo-cytoplasmic accumulating β-catenin cells (green, white arrowhead) which co-express the pituitary stem cell marker Sox2. Sox9+ve cells and proliferating cells (pHH3+ve cells) are found around the β-catenin+ve cell clusters (white arrows). Images are representative of 3 embryos per genotype. Abbreviations: AL, anterior lobe; IL, intermediate lobe; IHC, immuno-histochemistry; MZ, marginal zone; PL, posterior lobe; Tam, tamoxifen. Scale bars in **b** and **c** represent 50 μm.

Interestingly, Tam-induced *Sox2*^*CreERT2/+*^*;Apc*^*LoxP15/15*^ embryos exhibited hyperplasia of the anterior pituitary gland with the presence of clusters of accumulating nucleo-cytoplasmic β-catenin (**Fig. 6 b-c**). *In situ* hybridisation on consecutive histological sections revealed β-catenin+ve clusters expressing *Axin2* and the senescence marker *p21* (**Fig. 6b**) similar to previously discussed *Apc*-driven models (**Fig. 2 & Fig.5**). These clusters are Sox2+ve, have low proliferating capacity (negative for pHH3) and are surrounded by Sox9+ve cells (**Fig. 6c**) in keeping with aCPs^6,7,19^.

These results indicate that the Sox2+ve pituitary stem cells bearing bi-allelic loss of the tumour suppressor *Apc* (*Sox2*^*CreERT2/+*^*;Apc*^*LoxP15/15*^) are sufficient to initiate aCP tumour formation resulting in clusters of accumulating β-catenin that express diagnostic hallmarks of aCPs.

### *Apc*-driven cell clusters with nuclear β-catenin translocation undergo SASP and secrete inflammatory mediators, chemokines, cytokines, keratins and developmental growth factors

β-catenin-mutated+ve aCPs are characterised by whorl-like clusters of accumulating nucleo-cytoplasmic β-catenin which undergo SASP^7,18^ and act as hubs of secreting factors into the tumour microenvironment^7,18,19^. To investigate whether *Apc*-driven tumours contain clusters that act as signalling hubs, and to compare their molecular signatures to human β-catenin-mutated+ve aCPs, we performed transcriptomic analyses from isolated cell clusters from our *Cre:Apc*^*LoxP15/15*^ model at embryonic stages aiming to identify early tumour transcriptional signatures. To isolate β-catenin+ve cell clusters we utilised the transgenic line *Tcf/Lef-1:H2B-GFP* as a readout of activated Wnt-pathway^51^. The *Tcf/Lef-1:H2B-GFP* transgenic line (hereafter referred to as *Lef-1:GFP* for simplicity) harbours a knock-in of the Wnt/β-catenin signalling reporter construct containing 6 *Tcf/Lef1* response elements that drive the expression of GFP enabling fluorescent marking of Wnt-responsive cells. This transgenic line has been shown to accurately recapitulate the known pattern of Wnt-responsive cells and, therefore, represents an accurate read-out of activated Wnt/β-catenin canonical pathway^51^. We crossed our model *Cre:Apc*^*LoxP15/15*^ to the *Lef-1:GFP* reporter strain to generate *Cre:Apc*^*LoxP15/15*^*;Lef-1:GFP* compound mutants and assessed if the β-catenin+ve clusters expressed GFP (**Fig. 7 a-b**).

**Fig 7.**
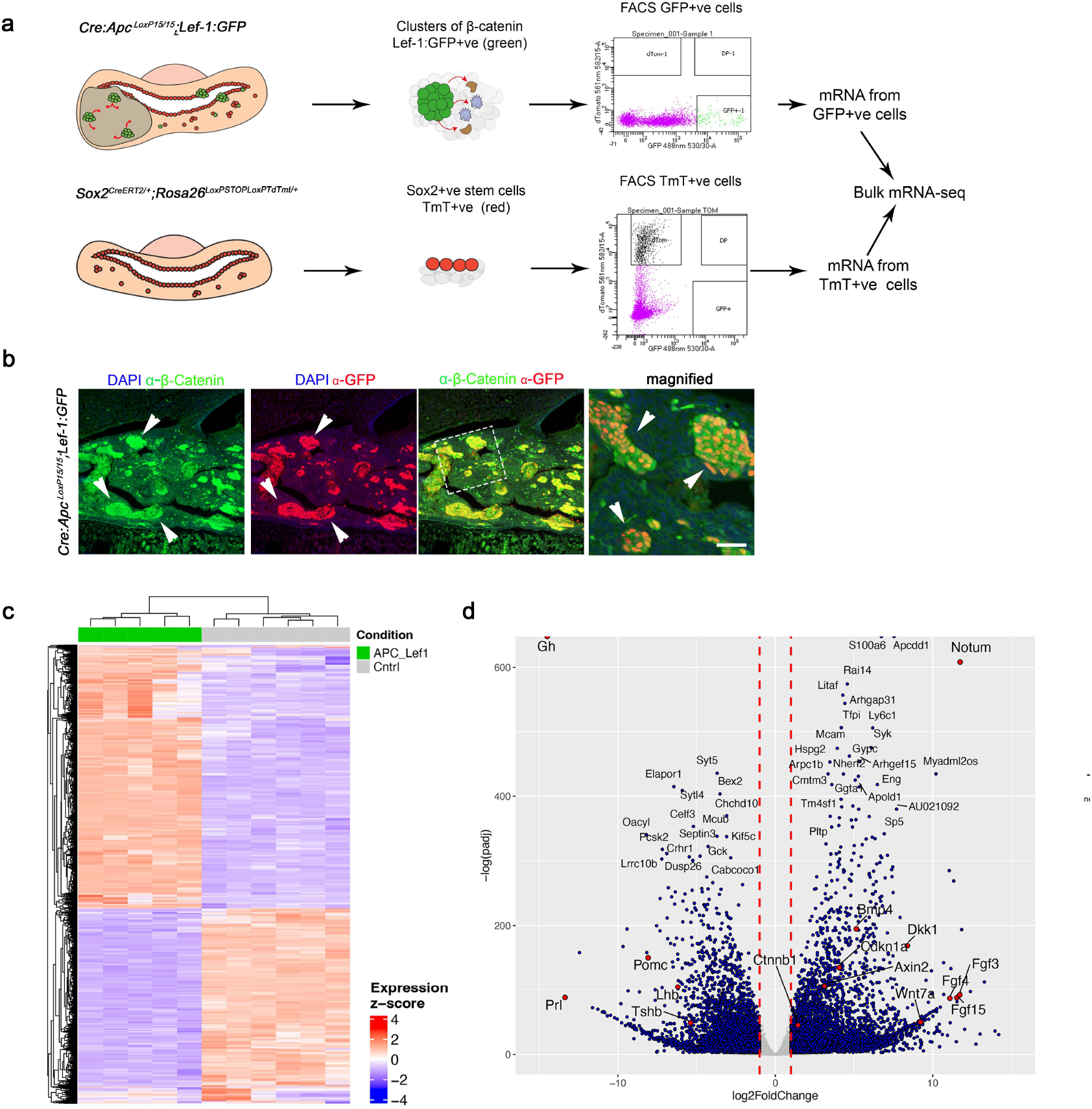
Transcriptomic analyses of early *Apc*-driven β-catenin+ve GFP+ve cell clusters show transcriptional signatures involved in inflammation, developmental growth factors and SASP. **a**) Diagram representing experimental design: mRNA sequencing from β-catenin+ve clusters from *Apc*-driven tumours (*Cre:Apc*^*LoxP15/15*^*;Lef-1:GFP*) was performed from the FAC-sorted GFP+ve fraction cells (β-catenin+ve GFP+ve cell clusters) and compared to Sox2+ve stem cells from Tam-induced *Sox2*^*CreERT2/+*^*;Rosa26*^*TdTomato*^ (Tomato+ve fraction cells). **b**) Double immunofluorescence against β-catenin (green) and α-GFP (red) revealed that all the nucleo-cytoplasmic β-catenin+ve cell clusters (white arrows) express GFP under the *Lef1* reporter sequence of the *Cre:Apc*^*LoxP15/15*^*;Lef-1:GFP*. **c**) Hierarchical clustering of the top 5000 most variable genes separates the cell population’s transcriptome from the *Apc*-driven β-catenin+ve GFP+ve cell clusters from control *Sox2*^*CreERT2/+*^*;Rosa26*^*TdTomato/+*^. **d**) Volcano plot showing differential gene expression between the β-catenin+ve GFP+ve cell clusters and control Sox2+ve pituitary stem cells. Highlighted in red are key upregulated genes such as the negative regulators of Wnt/β-catenin pathway *Notum, Dkk1;* the downstream effectors of activated Wnt/β-catenin pathway such as *Axin2, Sp5, Wnt7a* indicating overactivation of the canonical Wnt/β-catenin pathways; the developmental secreted factors *Fgf3, Fgf4, Fgf15* and *Bmp4* are found upregulated in addition to the senescence-associated factor *p21 (Cdkn1a)*. In line with aCPs being non-secreting tumours, pituitary terminal differentiation factors such as *Gh, Pomc, Lhb, Tsh* and *Prl* were downregulated. Abbreviations: *Gh*, growth hormone; FACs, fluorescent activating cell sorting; *Lhb*, luteinising hormone; Prl, prolactin; Tsh, thyroid-stimulating hormone. Scale bar in b represents 40 μm and the images are representative of 4/5 mutant embryos.

Double immunofluorescence against β-catenin and GFP revealed that all β-catenin+ve clusters, which over-activate Wnt-downstream targets, expressed GFP (**Fig. 7b**). Moreover, no GFP+ve cells were observed outside of these cell clusters, enabling accurate and specific fluorescent labelling of the β-catenin+ve cell clusters using GFP and the ability to perform purification using fluorescence activated cell sorting (FACs) (**Fig. 7a**). Tumours were dissociated to single cells and the GFP+ve fraction (β-catenin+ve GFP+ve cell clusters) was FAC-sorted before mRNA was extracted, purified and submitted for transcriptomic analyses. The mRNA from the GFP+ve cell fraction was compared to mRNA isolated from Wt pituitary stem cells from the *Sox2*^*CreERT2/+*^*;Rosa26*^*LoxPSTOPLoxPTdTmT*^ which allows for FAC-sorting of Sox2+ve cells expressing the red fluorescent protein TdTomato (or TdTmt for simplicity) (**Fig. 7a**). Hierarchical clustering of the most variable expressed genes distinguished transcriptional differences between the β-catenin+ve-GFP+ve cluster cells from that of the Sox2+ve TdTmt+ve cell population (**Fig. 7c**). Volcano plot analysis of differentially expressed genes revealed that *Apc*-driven clusters (β-catenin+ve GFP+ve) downregulate pituitary hormones such *Gh, Tsh, Pomc, Prl* and *Lh* (**Fig. 7d**) in accordance with the non-secreting nature of aCPs and is in keeping with our previous immunostaining analyses (**Fig. 1b & Fig. 4d**). Upregulated genes indicated that β-catenin+ve GFP+ve cell clusters activate the Wnt/β-catenin canonical pathway: with increase in the Wnt-downstream targets *Axin2, Lef1, Sp5, Tcf4*; and the secreted Wnt ligands: *Wnt7a, Wnt6, Wnt10a, Wnt9b, Wnt11, Wnt16* and their receptors and co-receptors *Fzd4, Fzd10, Lgr6*. Moreover, the secreted Wnt antagonists *Notum, Dkk1, Dkk2, Dkk4, Dkk3* and *Wif1* were found to be highly upregulated in the *Apc-*driven β-catenin+ve GFP+ve cell clusters (**Supplementary Fig. 3a**). We identified that genes such as the secreted fibroblast growth factor (*Fgfs*): *Fgf3, Fgf15, Fgf4, Fgf8, Fgf20* and *Fgf17*, the bone morphogenic factors: *Bmp4, Bmp8a, Bmp2, Bmp5, Bmp15* and *Shh* were found highly upregulated (**Supplementary Fig. 3b-c**). Keratins and keratin-associated proteins were highly upregulated by the *Apc*-driven β-catenin+ve GFP+ve clusters, which may contribute to the large deposits of wet keratin observed in our *Apc*-mutant model (**Fig. 4 e, Supplementary Fig. 4 a-b**). We identified that the GFP+ve β-catenin+ve clusters secrete a large number of interleukins (*Il6,Ilb, Il2b, Il36b*,*Il10, Il16, Il15*) and chemokines (*Ccl2, Ccl3, Ccl4, Ccl5, Ccl7, Ccl11, Cxcl9, Cxcl15, Cxcl3, Cxcl1, Cxcl11*) which have been reported to have proinflammatory, chemotactic, immunosuppressor and pro-angiogenic functions during tumour development and are the main components of SASP^52-55^ (**Supplementary Fig. 5 a-c**). In agreement with SASP, we identified that the SASP-associated factor p21 (*Cdkn1a*) was highly upregulated in the *Apc*-driven β-catenin+ve GFP+ve cell clusters which was also confirmed by *in situ* hybridisation (**Fig. 5d**). Some of the above-upregulated genes were found overexpressed in our *Apc*-model by *in situ* hybridisation in a cell cluster fashion (**Fig. 5d**) and mRNA-seq data was validated by RT-qPCR using mRNA isolated from *Apc-*driven GFP+ve β-catenin+ve cell clusters (**Supplementary Fig. 4c**). Unsupervised gene set enrichment analyses (GSEA) identified pathways upregulated by the *Apc*-driven β-catenin+ve GFP+ve cell clusters, such as positive regulation of inflammation, enrichment of IL2-Stat5 and IL6-Jak-Stat3 signalling pathways. These pathways are important in immunomodulation, inflammation and angiogenesis^53-55^ (**Fig. 8a**), reinforcing the hypothesis that β-catenin+ve GFP+ve clusters act as paracrine secreting hubs. Genes involved in positive regulation of TNF-alpha as well as KRAS and p53 pathways were found upregulated in cluster cells (**Fig. 8a**). We then compared our mRNA-seq transcriptomic data from our *Apc*-driven β-catenin+ve GFP+ve clusters to publicly available bulk mRNA-seq transcriptomics data sets, including human β-catenin-mutated+ve aCP bulk mRNA-seq data^19^ and data from murine models that express a degradation resistant form of β-catenin within Sox2+ve pituitary stem cells (*Sox2*^*CreERT2/+*^*;Ctnnb1*^*loxP(Exon3)/+*^)^18^ (**Fig. 8 b-c**). Gene Ontology (GO) analyses identified common upregulated pathways between our murine *Apc*-driven model and human β-catenin-mutated+ve aCPs, such as keratinisation, with a high number of keratin and keratin-associated proteins found upregulated (**Fig. 8c & Supplementary Fig. 4 a-b**), inflammatory response, neutrophil chemotaxis, IL2, IL6 and positive regulators of angiogenesis. These pathways are all linked to SASP mechanisms known to occur in β-catenin-mutated+ve aCPs^18-20,49^. One distinction identified in our transcriptomic analyses is that the *Apc*-driven β-catenin+ve GFP+ve clusters do not upregulate the senescence-associated factor p16 (Cdkn2a), in contrast to the p16 upregulation reported in some human and murine β-catenin-mutated+ve aCP clusters^18,19^. Together our data show that *Apc*-driven β-catenin+ve GFP+ve clusters are transcriptionally analogous, despite some differences, to previously described β-catenin-mutated+ve aCP cell clusters. These clusters exhibit SASP activity and function as secretory hubs within the tumour microenvironment in both murine^7,18^ and human^19,49^ aCPs.

**Fig 8.**
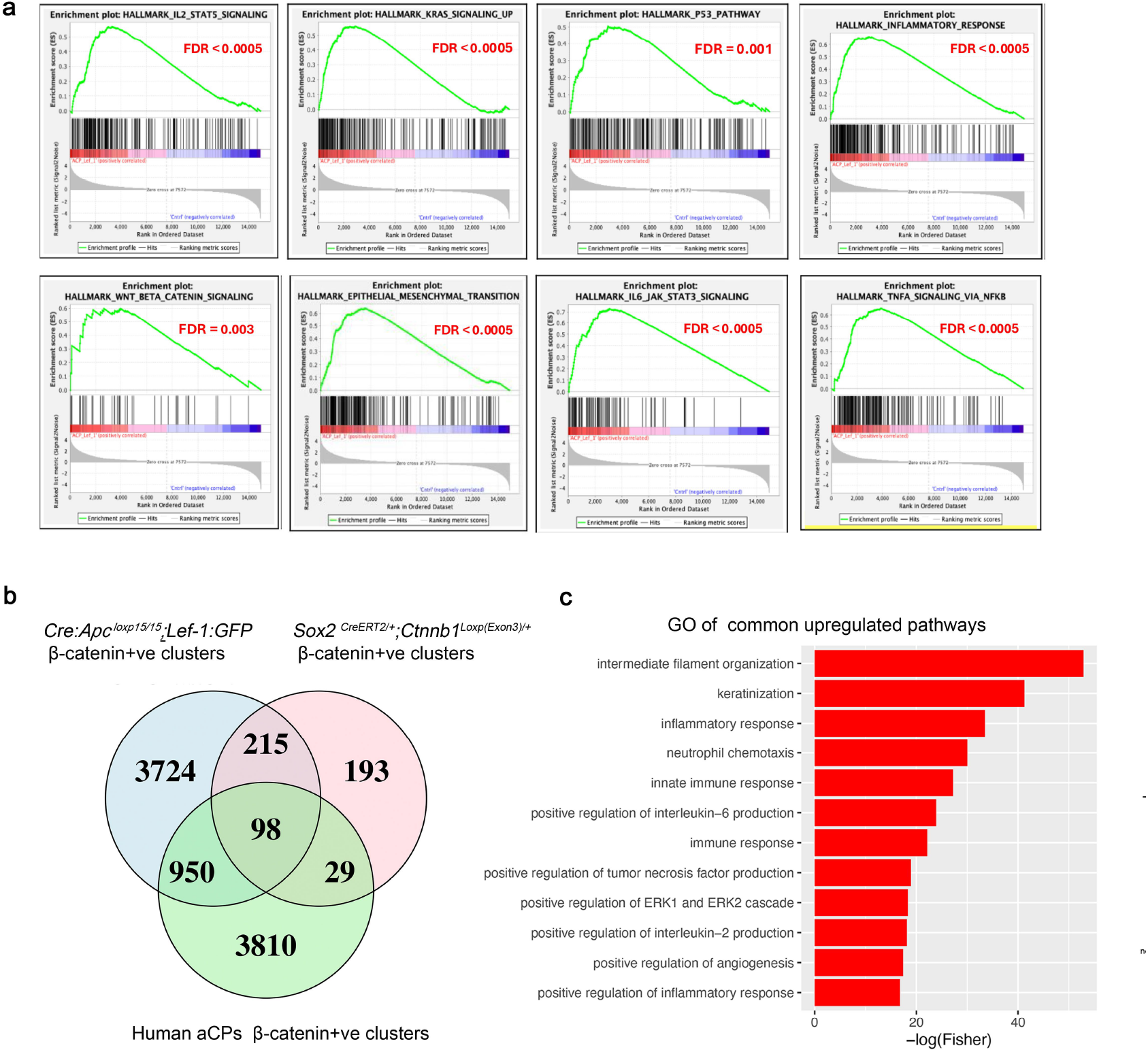
Transcriptomic analysis shows that *Apc*-driven tumours recapitulate human aCPs. **a**) Unsupervised Gene Set Enrichment analyses confirm that the β-catenin+ve GFP+ve cell clusters over-activate the Wnt/β-catenin canonical pathway with statistically significant upregulation of IL-2-Stat5, IL6-Stat3 and TNF-α signalling pathways involved in inflammatory response and SASP. The KRAS and p53 pathways were also found upregulated by the β-catenin+ve GFP+ve cell clusters, as were genes involved in epithelial-to-mesenchymal transition. **b**) Venn diagram comparing differential gene expression between *Cre:Apc*^*LoxP15/15*^;*Lef-1:GFP* derived clusters and clusters from β-catenin-mutated+ve model (*Sox2*^*CreERT2/+*^*;Ctnnb1*^*Loxp(Exon3*)/+^) and from human aCPs reveals higher number of genes in common between human aCP clusters and *Apc*-driven β-catenin+ve GFP+ve clusters (950 genes in common). **c**) Gene Ontology enrichment indicates upregulated pathways in both human aCPs, β-catenin-mutated+ve clusters and our model. These upregulated transcriptional signatures are important in aCP development and include pathways involved with keratinization, inflammasome: neutrophil chemotaxis, interleukin 6 and 2, angiogenesis, the innate immune response, positive regulation of inflammatory response and MAPK pathways. This indicates that the *Apc*-driven β-catenin+ve GFP+ve clusters are analogous to their β-catenin-mutated+ve aCPs. Abbreviations: GO, Gene Ontology.

## Discussion

Previous genetic studies in mice^6,7^ and humans^46,56^ have demonstrated that somatic mutations leading to a non-degradable form of β-catenin are the main driver in a substantial proportion of aCPs. However, the exact prevalence of β-catenin mutations in aCPs varies considerably between published studies^10-14^ and a significant fraction of aCPs lack a clearly defined driving mutation. Recently, two studies presented an association between mutations in *APC* leading to FAP in three patients reported to carry germline truncated mutations in *APC*^31,32^. A further possible association between aCPs and either FAP^26^, or *APC*-pathogenic syndromes such as Gardner^27^ and Turcot syndromes^28^ has been reported in 8 patients developing aCPs^34-43^. However, the lack of full genomic sequencing data for these patients’ tumours has not resolved whether mutations in *APC* had a direct role in the causation and formation of their aCPs. In this paper, we use three transgenic models to unequivocally show that disruption of *Apc* in the murine pituitary progenitors/stem cells is a main driver of aCPs independent of mutations in β-catenin. Firstly, we show that biallelic loss of *Apc exon15 (Cre:Apc*^*LoxP15/15*^) leads to severe intrauterine aCP tumours, which are not compatible with postnatal life. Importantly, cases of severe intrauterine aCPs have been reported by both our group^8^ and others^57,58^, indicating that aCPs can present at any stage of life. The severity of the phenotype with lack of all hormone-producing cells, leads to early perinatal lethality in this model. We developed a second model by generating a hypomorphic allele for *Apc* in which 7.5 % of wild type *Apc* mRNA is still produced (*Cre:Apc*^*LoxP1-15/15*^) leading to 100% penetrant postnatal aCPs. In this *Apc*-hypomorphic allele, postnatal tumours develop that have all the histopathological and molecular hallmarks of human aCPs, including large deposits of wet keratin, which were not observed in previously published β-catenin-mutated+ve aCP murine models^6,18^. Importantly, this new *Apc*-driven aCP model, faithfully recapitulates one of the most important aCP-disease comorbidities, tumour-induced morbid obesity, which was not seen in the previous murine model of aCPs driven by oncogenic β-catenin (*Hesx1*^*Cre/+*^*;Ctnnb1*^*Loxp(Exon3)/+*^)^6,18^. Thus, our model represents a robust genetic tool to study aCP tumour-induced hypothalamic obesity, a major complication seen in aCP patients^2,59-62^. It is worth noting that in our *Apc*-driven aCPs (*Cre:Apc*^*LoxP1- 15/15*^), we observed gender differences in tumour progression with females living longer than males. Whether this reflects a difference in tumour immune microenvironment between genders, as seen in models of *Wnt*-driven adrenocortical carcinoma (ACCs)^63^ d will require further investigation. However, human epidemiological studies from large aCP patient registries do not show gender differences in disease progression for β-catenin-mutated+ve aCPs^2,5,64,65^. Our mouse genetic study provides further information on the role of Apc in tumorigenesis and argues against a dominant negative effect of truncated *Apc* which has been hypothesised in other cancers^66,67^. Our hypomorphic model of aCPs (*Cre:Apc*^*LoxP1-15/15*^) expresses 7.5% of wild type *Apc*. This is sufficient to rescue the early perinatal lethality observed with bi-allelic loss of *Apc exon15*, which is typically associated with more aggressive and larger gestationally derived tumours. Thus, our data show that small amounts of Wt *Apc* mRNA is sufficient to ameliorate the severe phenotype arguing against a dominant negative role of truncated *Apc exon15*.

β-catenin-mutated+ve aCPs have been shown to originate from pituitary progenitors/stem cells that express the stem cell marker Sox2^7^. Using a transgenic *Sox2*^*CreERT2/+*^ to delete *Apc exon15*, we show that biallelic loss of *Apc exon15* in the Sox2+ve pituitary stem cells is sufficient to produce early aCP lesions indicating that these Apc-tumours arise from progenitor/stem cells of the pituitary gland. Some aCPs are characterised histologically by the presence of nucleo-cytoplasmic β-catenin+ve cell clusters that are a diagnostic hallmark of aCPs^6,11,17,46^. These clusters have been shown to initiate tumour formation and have been studied extensively in β-catenin-mutated+ve aCPs both in mice^6,17,18^ and humans^17,19,68^. These nucleo-cytoplasmic β-catenin+ve cell clusters exhibit SASP activity, secreting proinflammatory mediators, chemokines, cytokines and growth factors that collectively alter the tumour microenvironment. Our transcriptomic analyses of *Apc*-driven clusters revealed that, indeed, these tumour-initiating cells undergo SASP and act as secretory hubs to the tumour microenvironment. We identified key secreted SASP factors such as proinflammatory mediators, abundant cytokines and chemokines, interleukins and angiogenesis factors, all of which have been associated with SASP in various tumours^53-55^. These clusters secrete numerous downstream targets of the Wnt canonical pathway and developmental growth factors such as Fgfs, Bmps and Shh, which may influence the tumour micro-environment. Indeed, a recent study has shown that in β-catenin-mutated+ve aCPs, clusters can modify tumour-associated macrophages from a M1 (anti-tumorigenic) to M2 (pro-tumorigenic) phenotype in co-culture experiments^68^, an effect that could be similar within *Apc-*driven tumours. Although, the *Apc-* driven β-catenin clusters seem largely analogous to their β-catenin-mutated+ve counterparts in humans, some transcriptional differences were noted that could confer specific targeted treatment. Compared to other previously published murine aCP models^6,18^, we identify that the *Apc*-driven clusters express a large number of keratins and keratin-associated proteins which contribute to the large wet-keratin deposits that parallel human aCPs which are absent in other published aCP murine models. Another difference within our *Apc*-driven tumour model is the mechanism of SASP, which relies on the p21/p53 pathway rather than Cdkn2a (p16), differing from previously published human and murine data on β-catenin-mutated+ve aCPs^17-19,49^. Whether these transcriptional differences are characteristic of all APC-mutated+ve tumours requires additional studies in more aCP samples to enable stratification of transcriptional signatures based on the aCP-mutation specific subtype. Our data call for in-depth genomic studies to identify the overall prevalence of *APC*-mutations in larger cohorts of patients using high depth of reading sequencing techniques, which will allow for detection of low allelic frequency mutations. These studies will lead to genotype/clinical phenotype correlation in patients without mutations in oncogenic β-catenin and identify whether their clinical trajectories differ i.e. number of recurrences and/or hypothalamic involvement. Importantly, our murine genetic study shows that dysfunction of *Apc* is a main driver of aCPs independent of mutations in β-catenin and hence APC-mutated+ve tumours represent a novel genetic subtype of aCPs. Since our data demonstrate the causal effect of Apc-dysfunction in the development of aCPs, this highlights the need for genetic testing and increased surveillance of patients with FAP or germline *APC*-pathogenic syndromes for the development of aCPs. Furthermore, it warrants the inclusion of aCPs as part of the broader spectrum of tumours associated with these syndromic conditions. Equally, patients that develop aCPs with β-catenin-mutated-negative tumours should be tested for underlying germline mutations within *APC* as a possible cause of FAP or APC-related syndromes such as Gardner and Turcot syndromes.

## Methods

### Animal

All experiments were conducted under regulations, licenses and local ethical review (QM-AWERB Ethical Committee) following the UK Home Office Animals (Scientific Procedures) Act 1986. The transgenic lines *Rosa26*^*CAGLoxpSTOPLoxpTdTomato*^ (stock #007905)^69^, *ApcLoxp(Exon15)/Loxp(Exon15)* (stock #029275)^43^, *ApcLoxp(Exon1-15)/Loxp(Exon1-15)* (stock #009045)^48^, *Tcf/Lef-1:H2B-GFP* (stock #032838)^51^ and the *Sox2*^*CreERT2/+*^ (stock #017593)^50^ were obtained from the JAX-lab and have been previously described. The *Prop1:Cre* transgenic line^44,45^ was kindly provided by Shannon Davis and Sally Camper. Animals were kept in 12 h light/12 h dark cycle, with a constant supply of food and water, temperatures of 65–75°F (∼18–23°C) with 40–60% humidity. Kaplan Meier survival curves were plotted using humane end points as determined in line with UK Home Office regulations regarding use of mice in research. Mice were humanly culled when health deterioration was assessed to be irreversible. Tamoxifen induction of the Cre recombinase in embryos (*Sox2*^*CreERT2/+;*^*Apc*^*LoxP15/15*^) during pregnancy was achieved by 3 consecutive daily injections of 0.1 mg/g of mouse at e12.5, e13.5, e14.5 days of gestation and embryos were dissected at e18.5.

### Immunohistochemistry, immunofluorescence, and *in situ* hybridisation

Immunostaining was performed as previously described^70-72^ in short: sections were deparaffinised followed by rehydration through decreasing ethanol dilutions. Heat-induced antigen retrieval was performed with a microwave in 10 mM sodium citrate buffer (pH 6). Samples were left to cool and incubated for 1 h in blocking buffer [1 PBS, 0.1% Triton X-100, 5% Normal Goat Serum (Vector Laboratories)]. Antibodies, their concentration and source are listed in **Supplementary Table 1**. Immunohistochemistry (IHC) staining was achieved using DAB Peroxidase Substrate Kit (Vector Laboratories; SK-4100). The colorimetric reaction was stopped with water and sections were counterstained using haematoxylin (Sigma-Aldrich). For immunofluorescence (IF), conjugated secondary antibodies Alexa Fluor 568 or 488 were used, or a biotinylated secondary followed by streptavidin (**Supplementary Table 1**). Sections were mounted with VECTASHIELD containing DAPI (Vector Laboratories). Images were acquired with a Leica microscope. Figures were generated with Adobe Photoshop CS9. The mitotic index (MI) is the percentage of pHH3-positive cells compared to total number of cells (average counts from three different sections, separated approximately by 100 μm, per embryo/pituitary with a minimum of *n* = 3–4 per genotype and stage). *In situ* hybridisation was performed by adapting the protocol from^73^ and described before in^72-75^. In short, slides were deparaffinised, rehydrated and fixed with 4% PFA. Slides were incubated with proteinase K, followed by a second fixation with 4% PFA and finally incubated with 0.1 M triethanolamine and 0.1% acetic anhydride (Sigma). Hybridisation was achieved by an overnight incubation with 100 ng of the digoxigenin-labelled probe at 65°C. Sections were washed in 0.1 M Tris-HCl Buffer (pH = 7.5) followed by an overnight incubation at 4 °C with anti-Dig antibody (Sigma-Aldrich). Signal detection was achieved by colorimetric reaction using 4-nitro blue tetrazolium chloride solution (NBT; Sigma-Aldrich) and 5-bromo-4-chloro-3-indolyl phosphate disodium salt (BCIP; Sigma-Aldrich). The digoxigenin-labelled antisense probes *p21, Axin2, Shh, Fgf8, Fgf3, Wnt6* and *Wnt7a* were generated from plasmids containing either a portion or full-length cDNA of each respective gene, and plasmids were obtained from Source Bioscience, or gifts from Sally Camper, Andreas Kispert and Peter Gruss and published before^76-78^. Pituitary volume quantification was performed using H&E stained pituitaries. Four sections in every 16 were counted for estimation. Microscope images were taken at 5x magnification and analysed using image J software. Estimation of pituitary volume was calculated by multiplying the area of the pituitary by the thickness of each section (7µm) and correcting for the remaining sections in the series [Pituitary volume (µm^3^) = Measured area(µm^2^) x 0.7(µm) x 4]. Comparative analyses of pituitary volumes at each embryonic stage from at least 3 different embryos per genotype from different litters were made using Student’s T-Test.

### Fluorescent activated cell sorting (FACs)

To FAC-sort pituitary cell populations, pituitary single cell suspension was achieved by enzymatic treatment using 250 µL enzyme mix [0.05% Collagenase Type II (LorneLaboratories Ltd), 50 µg/ml DNaseI (Worthington), 2.5μg/ml Amphotericin B (Gibco), 0.1% v/v trypsin-EDTA solution 0.05% (Gibco) in Hank’s Balanced Salt Solution (HBSS, Gibco)] in an RNAse and DNAse free 1.5ml microcentrifuge tube (Axygen). Samples were incubated for 1 hour at 37°C under constant agitation at 500 rpm. Pituitaries were mechanically dissociated by repeated pipetting. Dead cells were stained with 500 µL of 1/5000 sterile DAPI solution incubated under gentle agitation for 3 minutes. Samples were centrifuged and cells were resuspended in 500 µL FACS buffer [25 mMHEPES (Gibco), 1% FCS (Sigma-Aldrich) in sterile PBS] by repeated pipetting and transferred to a flow cytometry tube. For mRNA purification, cells were FAC-sorted directly in 750 µL of TRIzolTM-LS (Invitrogen). Cells were sorted using a BD FACs Aria II sorter (BD bioscience) using a A100 µm nozzle. Side scatter (SSCr) area and forward scatter (FSC) area were compared to ensure cellular matter was identified and sorted. SSCr width and FSC area were then compared to ensure only single cells were sorted. A 405 nm laser was used to sort DAPI+ve and DAPI-ve cells. A 561 nm laser was used to sort tdTomato+ve and tdTomato-ve cell populations. A 488 nm laser was used to sort GFP+ve and GFP-ve cells. Sorted cells were immediately snap frozen and stored at -80°C until mRNA purification.

### RNA purification, RT-qPCR for recombination efficiency and validation of mRNA-seq

RNA from FAC-sorted cells from *Sox2*^*CreERT2/+*^*;Rosa26*^*LoxpSTOPLoxPtdTmT*^, *Cre:Apc*^*LoxP15/15*^*;Lef-1:GFP* or stem cell cultures from *Cre:Apc*^*15/15*^ or Wt pituitaries was obtained using PureLink™ RNA Micro Kit (Invitrogen) following the manufacture’s protocols. Samples were submitted to quality control using an Agilent 2100 Bioanalyser and only samples with RNA integrity values ≥ 8.1 were kept for mRNA-seq analyses. To generate cDNA for RT-qPCR, 500 ng of RNA was incubated at room temperature for 15 minutes with 1 µL of DNAse I (Invitrogen), 1 µL 10x DNase I Reaction Buffer and UltraPure RNAse free water (Invitrogen) to a total volume of 10 µL. Reverse Transcription was performed using a High-Capacity cDNA Reverse Transcription Kit (Applied Biosystems) to produce cDNA according to the manufacturers protocol. Quantity and quality of the cDNA samples were analysed using Nanodrop ND-1000 spectrophotometer (Nanodrop) and quality of cDNA was assessed by determination of the 260/280 and 260/230 ratios. Gene expression levels were analysed by RT-qPCR using QuantiTect SYBR Green PCR Kit (Qiagen) according to the manufacturer’s protocol and analysed with Stratagene (Agilent Technologies). A comparative Ct method (2^−ΔΔCT2^ method) was used to compare the mRNA expression levels of genes of interest between cell populations normalised to GAPDH. RT-qPCR primers for *GAPDH, Apc exon15, Axin2, Wnt7a, Notum, Fgf3, Ccl2, Ccl3, Gh, Tsh, Pomc, Prl* are shown in **Supplementary Table 2**.

### Pituitary stem cells (PSCs) culture and proliferation assays

PSCs were cultured from Wt or *Cre:Apc*^*loxP15/1-15*^ mutant pituitaries and incubated for 2 h in enzyme mix [0.5% w/v Collagenase, 50 µg/ml DNase, 2.5 µg/ml Fungizone, trypsin 0.1% in Hank’s Balanced Salt Solution] and mechanically dissociated into single cells. In all, 10,000 cells/well were plated in 12-well plates and cultured in stem cell-promoting media [Ultraculture Medium (Lonza), supplemented with 5% FCS (Sigma), 1% penicillin/streptomycin (P/S: Fisher), 1% glutamax (Fisher), 20 ng/ml basic fibroblast growth factor (R&D) and 50 ng/ml cholera toxin (Sigma)]^72^. Media was changed every 48 h and cultures were maintained for 8 days. Cells were stained with Coomassie blue (Fisher), and either the number of colonies or cells per colony was counted.

### Statistics and reproducibility

Statistical analyses were performed using Prism9 software (GraphPad). The number of independent experiments and replicates (*n*) is indicated in each of the figure legends. Unless stated otherwise, at least three biologically independent replicates were performed for each panel, which came from at least three independent experiments. When appropriate, normalisation of the data was performed within each independent experiment.

### mRNA-seq and bioinformatic analyses

For library preparation mRNA enrichment for cell samples using the SMARTer ultralow input kit (Illumina), and 150bp paired-end sequencing was then performed on a NovaSeq6000 platform. Reads were quality trimmed using Trimgalore v0.6.5 and aligned to reference genome GRCm39 using STAR v2.7.9a^79^. Reads were counted using feature counts, and the ENSEMBL annotation gencode.vM33.annotation.gtf. Downstream analysis was performed in DESeq2^80^, with filtering of genes with less than one read count per million (cpm). Genes with a |logFC|>2 and p.adj <0.05 were considered significant. Gene Ontology (GO) enrichment was calculated on the most strongly upregulated genes (logFC>5) and downregulated (logFC< -5) genes separately using the TopGO v2.52.0 package. For unsupervised hierarchical clustering, the top 5000 variable genes (ranked by standard deviation) were used. For comparison to the BcatGOF model^7^ and human aCP^19^ published data raw fastqs were accessed from the European Nucleotide Archive (PRJEB18056) and analysed as above using GRCh38.release109 and ENSEMBL annotation Homo_sapiens.GRCh38.109.gtf for the human samples.

## Data availability

The authors declare that all the data supporting the findings herein are included in the article (or Supplementary materials) and available from the corresponding author (C.G.M.) upon reasonable request. All sequencing raw data files have been deposited to GEO under reference GSE282683 (https://www.ncbi.nlm.nih.gov/geo/query/acc.cgi?acc=GSE282683). The source data underlying graphs and supplementary materials are provided as a Source Data file. A reporting summary for this article is available as a Supplementary Information file.

## Acknowledgements

We apologise to the authors whose work we were unable to reference due to space constraints. The research is funded by Action Medical Research (GN 2272), Grants from Barts Charity (GN 417/2238 and MGU0551) and Sparks-GOS (V4323). R.T. and J.B. were funded by Medical Research Council, MRC-CRTF (MR/P018459/1) and (MR/S037896) grants respectively. LGC funded by PhD from Barts Charity (MGU0551). AR was funded by Newton-Bhabha Fund, Academy of Medical Sciences UK (NIFR8\1037).

## Author Contributions

CGM conceived the work, obtained funding, supervised research, performed experiments, analysed data, wrote the manuscript and dealt with all the editorial publishing process. JB, LGC, JN, AR, RT, JX, KCG, AG, PD performed experiments and analysed data. JN, CH, AR bioinformatic analyses. JB, RT & JN contributed to funding acquisition. SWD, provided the *Prop1:Cre* transgenic line. PD, MK supervised AR JB. FR provided neuropathological expertise. All authors helped with the editing and agreed with its content for publication.

