## Supplementary Figures and Tables for "Disruption of the Wnt-antagonist *Apc* in the pituitary stem cells drives the development of adamantinomatous craniopharyngioma"

### Supplementary material

**Supplementary Fig 1. Deletion of *Apc* exon1 to exon15 in the pituitary progenitors (*$Cre:Apc^{LoxP1-15/1-15}$*) results in hyperplasia of the pituitary gland but not tumour formation. a**) IHC of coronal sections through the pituitary gland and brain at e18.5 against GH, POMC, PRL and TSH of Wt (left column) or *$Cre:Apc^{LoxP1-15/1-15}$* mutants (right column) reveals mutant pituitaries exhibit hyperplasia with decreased GH hormone-producing cells. **b**) IHC against the proliferation marker pHH3 at e12.5, e15.5 and e18.5 reveals an increased number of pHH3+ve cells in *$Cre:Apc^{LoxP1-15/1-15}$* mutants compared to Wt pituitaries leading to a statistically significantly increased mitotic index (MI) **c**. **d**) RT-qPCR to amplify exon15 of Apc using Wt or FAC-sorted cells from *$Cre:Apc^{LoxP1-15/1-15}$*,*$Cre:Apc^{LoxP15/15}$* or the compound heterozygote *$Cre:Apc^{LoxP1-15/15}$*. Exon15 was fully recombined in the *$Cre:Apc^{LoxP15/15}$*, but exon 15 of *Apc* was detected in the *$Cre:Apc^{LoxP1-15/1-15}$* at 16.5% and 7.5% in the *$Cre:Apc^{LoxP1-15/15}$* indicating incomplete recombination of the *Apc LoxP exon 1-15* locus. *P* values in **c** were obtained using unpaired two-tailed Student's *T*-test and data represented as mean ± SEM, from 3 pituitaries per genotype. Images are representative of 4-5 embryos per genotype. Abbreviations: AL, anterior lobe; LH, luteinizing hormone; GH, growth hormone; H&E, haematoxylin and eosin; IHC, immuno-histochemistry; MZ, marginal zone; PL, posterior lobe; POMC, Pro-opiomelanocortin; PRL, prolactin; SB, sphenoid bone; TSH thyroid-stimulating hormone. Scale bars in **a** and **b** represent 100 μm.

**Supplementary Fig 2. The *$Cre;Apc^{LoxP15/15}$* mutant pituitaries exhibit larger nucleo-cytoplasmic β-catenin accumulating cell clusters compared to the *$Cre;Apc^{LoxP1-15/15}$* hypomorph**. **a**) Immunofluorescence against β-catenin (green) on coronal sections through the pituitary glands at e18.5 reveals that the *$Cre;Apc^{LoxP15/15}$* mutant pituitaries have larger nucleo-cytoplasmic β-catenin cell clusters compared to the *$Cre;Apc^{LoxP1-15/15}$* hypomorphic allele suggesting a milder phenotype. **b-c**) The number of clusters per pituitary were the same between genotypes, but volume of the β-catenin cell clusters was larger in *$Cre;Apc^{LoxP15/15}$* mutant pituitaries. Images are representative of 4 to 5 embryos per genotype. *P* values in **c** were obtained using unpaired two-tailed Student's T-test and data represented as mean ± SEM, from 3 pituitaries per genotype. Abbreviations: AL, anterior lobe; MZ, marginal zone; PL, posterior lobe; SB, sphenoid bone. Scale bars in **a** represent 100 μm.

**Supplementary Fig 3. The β-catenin+ve GFP+ve cell clusters from *$Cre:Apc^{LoxP15/15}$;Lef-1:GFP* tumours over activate the Wnt/β-catenin canonical, fibroblast growth factor (Fgfs) and bone morphogenetic (Bmps) pathways**. **a**) Graph representing $Log_2$ fold change of upregulated genes belonging to the Wnt/β-catenin canonical pathway. The Wnt-downstream targets *Axin2*, *Lef1*, *Sp5*, *Tcf4*; and the secreted ligands: *Wnt7a*, *Wnt6*, *Wnt10a*, *Wnt9b*, *Wnt2*, *Wnt16, Wnt11* and their receptors and co-receptors *Fzd4*, *Fzd10*, *Lrp4, Lgr6*. Moreover, increase in secreted Wnt antagonist *Notum, Dkk1, Wif1*, *Dkk4*, *Dkk2*, *Dkk3* were found to be highly upregulated in the *Apc*-driven β-catenin+ve GFP+ve cell clusters. **b**) Graph representing $Log_2$ fold change of upregulated genes belonging to the secreted fibroblast growth factor (*Fgfs*): *Fgf3*, *Fgf15*, *Fgf4*, *Fgf8, Fgf20* and *Fgf17* and the bone morphogenic factors (**c**): *Bmp4, Bmp8a*, *Bmp2*, *Bmp5*, *Bmp15* were found upregulated by the *Apc*-driven β-catenin+ve GFP+ve cell clusters.

**Supplementary Fig 4. The β-catenin+ve GFP+ve cell clusters from *$Cre:Apc^{LoxP15/15}$;Lef-1:GFP* tumours over express cytokeratin and cytokeratin associated proteins**. **a**) Graph representing $Log_2$ fold change of the most upregulated cytokeratins identified from the transcriptomic analyses. **b**) Graph representing $Log_2$ fold change of the most upregulated cytokeratin associated protein identified in the transcriptomic analyses. **c**) Validation of the *Apc*-driven β-catenin+ve GFP+ve cell clusters transcriptomic analyses by RT-qPCR of some of the upregulated genes (red) *Axin2* ($p<0.0001$), *Lef1* (*p*=0.0025), *Wnt7a* (*p*=0.0001), *Notum* (*p*=0.0024), *Fgf3* (*p*=0.0039); *Ccl2* (*p*=0.0043), *Ccl3* (*p*=0.0003) and downregulated genes (blue) *Gh* (*p*=0.0001), *Tsh* (*p*=0.0005), *Pomc* (*p*=0.0004), *Prl* (*p*=0.0001). *P* values in **c** were obtained using unpaired two-tailed Student's *T*-test and data represented as mean ± SEM, from 3-4 pituitaries per genotype.

**Supplementary Fig 5. The β-catenin+ve GFP+ve cell clusters from *Cre:Apc*$^{LoxP15/15}$*;Lef-1:GFP* tumours overexpress high levels of multiple cytokines, chemokines and interleukins**. **a**) Graph representing $Log_2$ fold change of the most upregulated chemokines. **b**) Graph representing $Log_2$ fold change of the most upregulated cytokines expressed by the β-catenin+ve GFP+ve cell clusters **c**). Graph representing $Log_2$ fold change of the most upregulated interleukins expressed by the β-catenin+ve GFP+ve cell clusters.

### Supplementary Figure 1

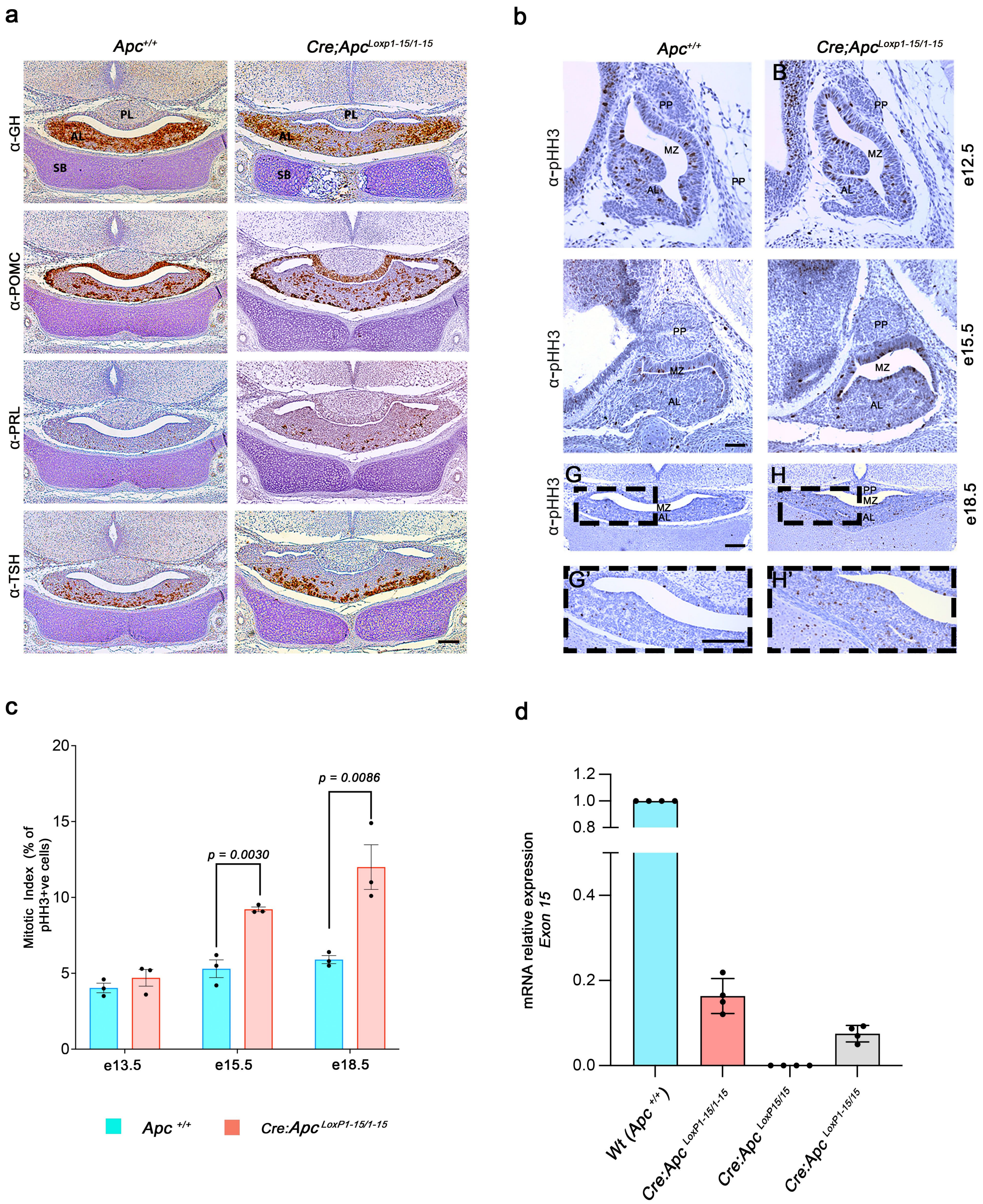

#### Supplementary Figure 2

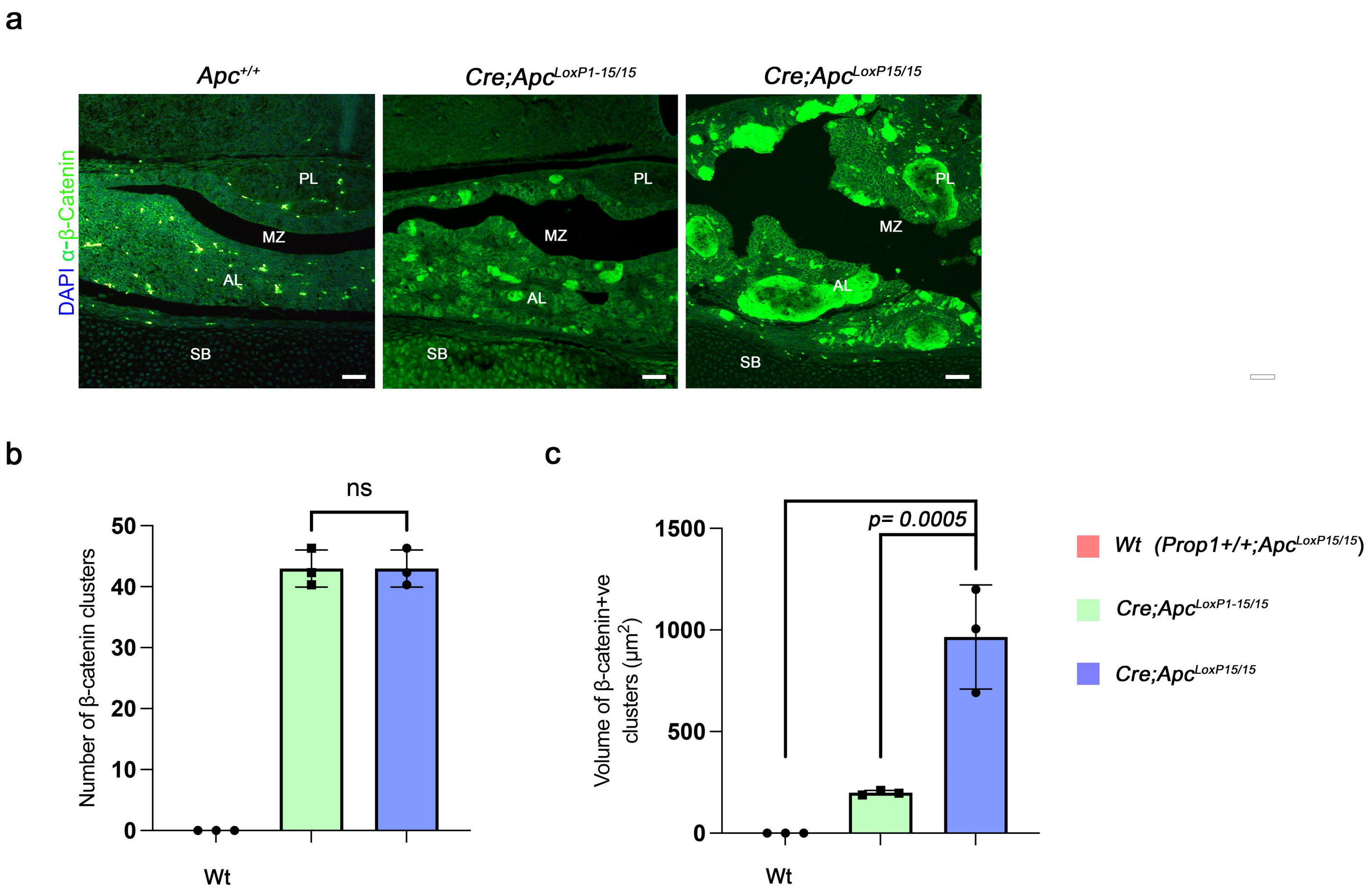

Suppplementary Figure 3

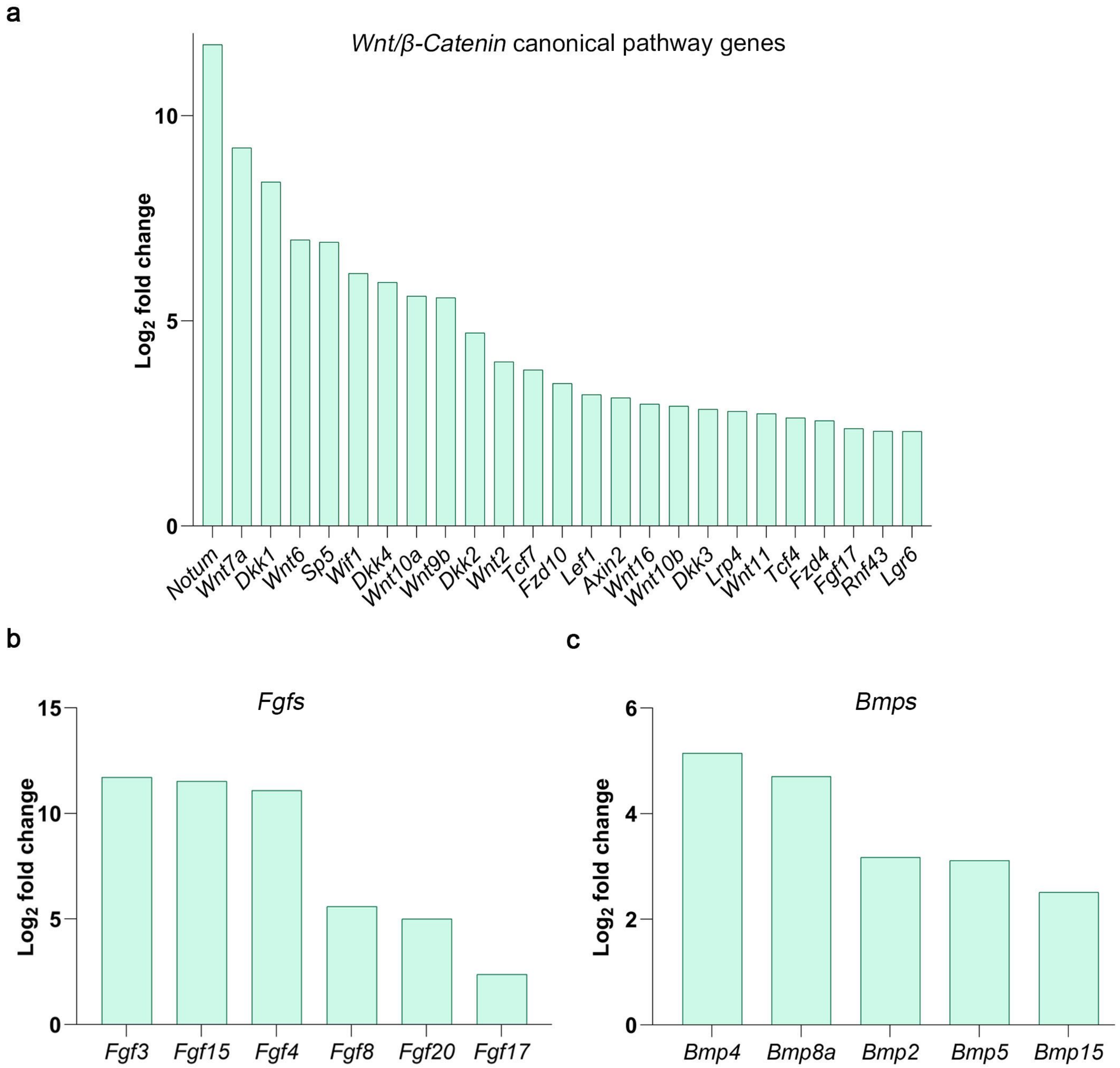

#### Supplementary Figure 4

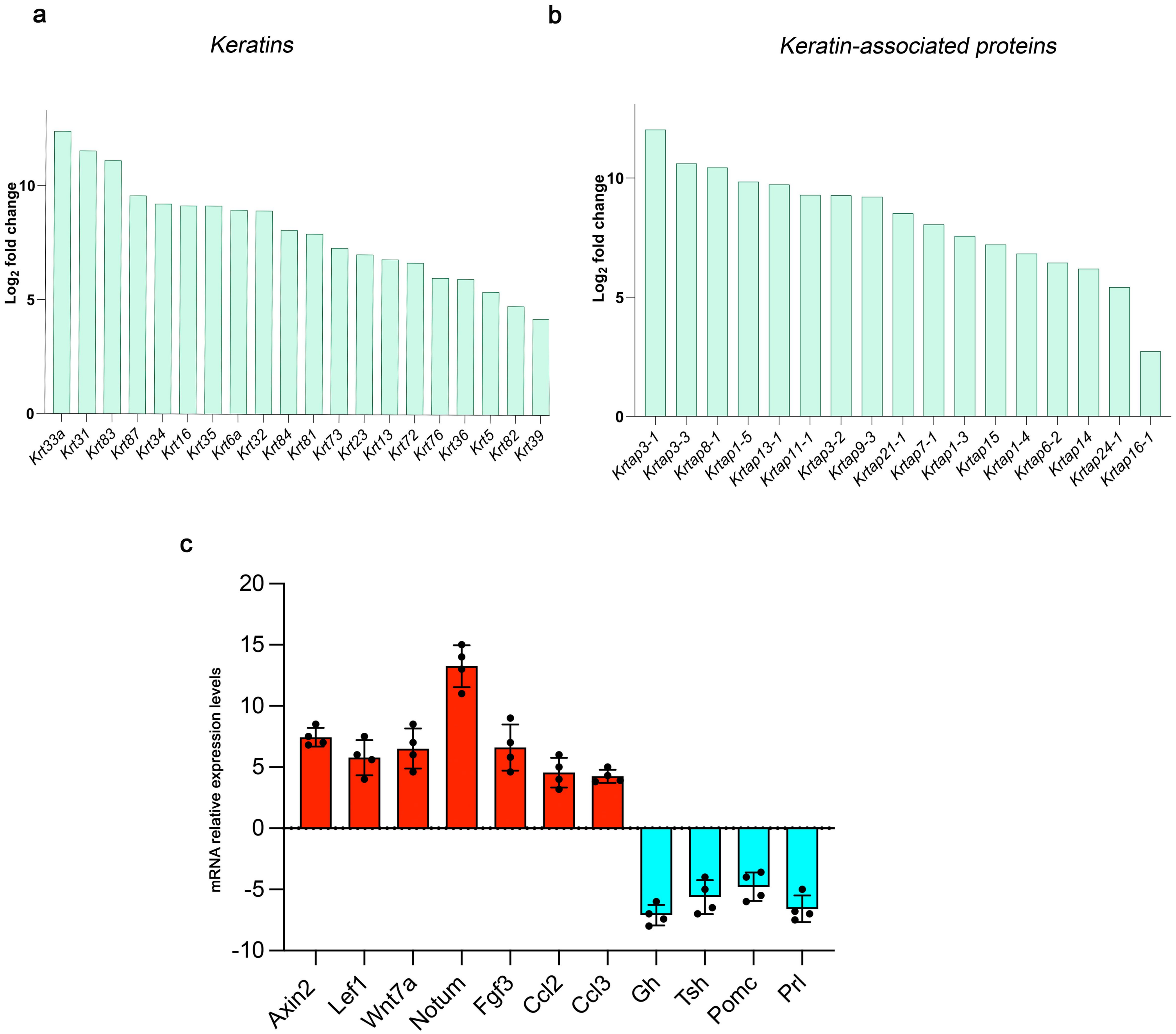

Supplemetary Figure 5

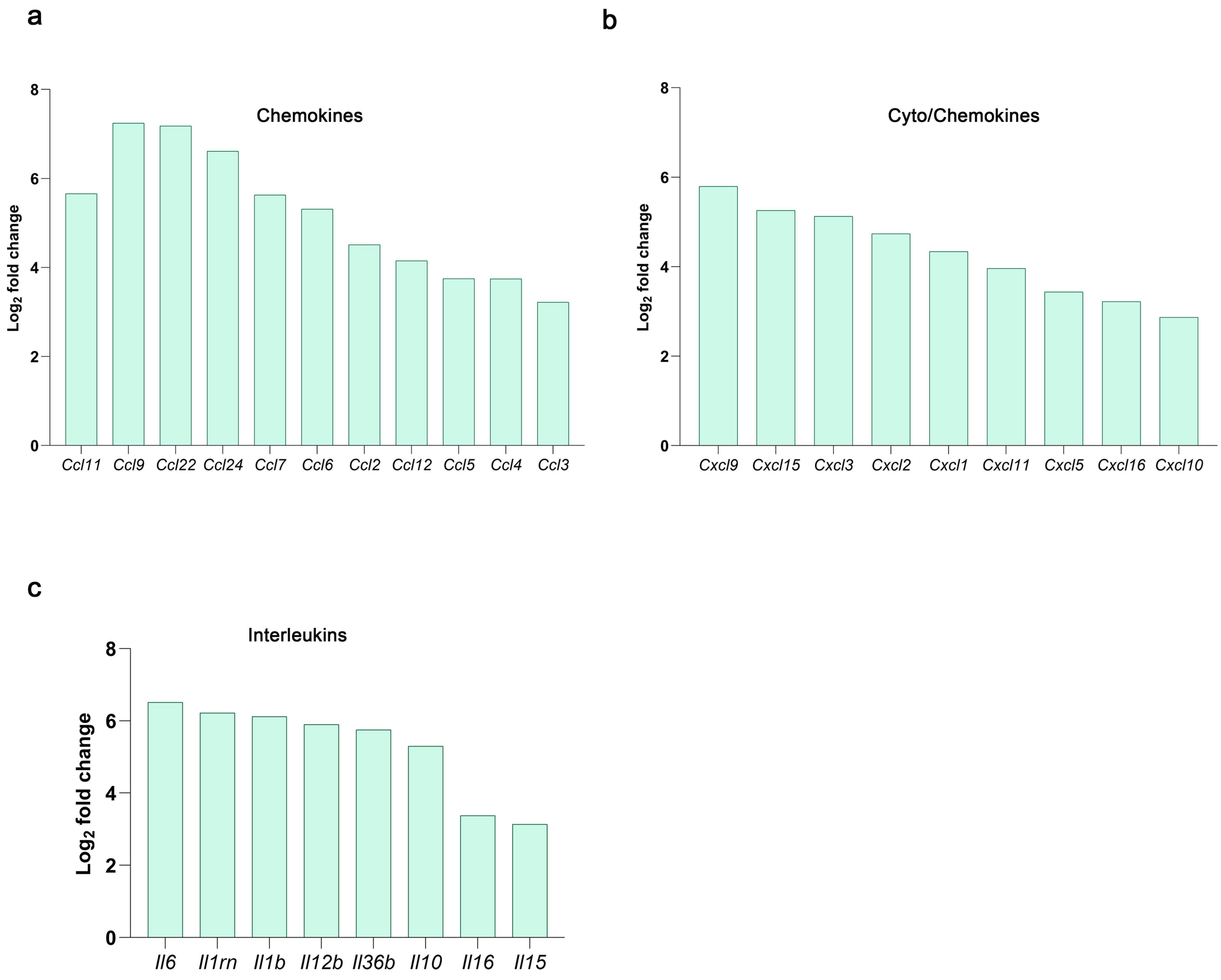

**Supplementary Table 1**. **Antibodies**

| Antibodies | Source |
|---|---|
| Rabbit α-phosphorylated-histone H3 ( α-pHH3) | 1:300 dilution, MILLIPORE, 06-570 |
| Rabbit α-PRL | 1:500 dilution, The National Hormone and Peptide Program (NHPP) Harbour-UCLA Medical Centre |
| Rabbit α-LH | 1:500 dilution, The National Hormone and Peptide Program (NHPP) Harbour-UCLA Medical Centre |
| Rabbit α-TSH | 1:500 dilution, The National Hormone and Peptide Program (NHPP) Harbour-UCLA Medical Centre |
| Rabbit α-GH | 1:500 dilution, The National Hormone and Peptide Program (NHPP) Harbour-UCLA Medical Centre |
| Rabbit α-Pomc | 1:500 dilution, The National Hormone and Peptide Program (NHPP) Harbour-UCLA Medical Centre |
| Rabbit α-GSU | 1:500 dilution, The National Hormone and Peptide Program (NHPP) Harbour-UCLA Medical Centre |
| Goat α-Sox2 | 1:200 dilution, Neuromics, GT15098 |
| Rabbit α-Sox9 | 1:500 dilution, Millipore, AB5535 |
| Mouse α-β-Catenin | 1:300 dilution, SIGMA, C7082 |
| Chicken α-GFP | 1:100 abcam ( ab13970) |
| biotinylated goat anti-rabbit antibody | 1:300 dilution, Vector Laboratories, BA-1000 |
| biotinylated horse anti-goat antibody | 1:200 dilution, Vector Laboratories, BA-9500 |
| donkey anti-goat antibody, Alexa Fluor 568 | 1:300, Invitrogen, A-11057 |
| goat anti-mouse antibody, Alexa Fluor 488 | 1:300, Invitrogen, A-11001 |

Table containing the primary and secondary antibodies used.

**Supplementary Table 2**. **Primers for RT-qPCR**

| Gene | Primer sequence |
|---|---|
| *Apc (exon 15)* | FW: ACGAAGACCACAGGCAAATC<br>REV: CCACAAAGTTCCACATGCATTAC |
| *Axin2* | FW: ACGAAGACCACAGGCAAATC<br>REV: CCACAAAGTTCCACATGCATTAC |
| *Lef1* | FW: ACTGTCAGGCGACACTTCCATG<br>REV: GTGCTCCTGTTTGACCTGAGGT |
| *Wnt7a* | FW: TTCGCCAAGGTCTTCGTGGATG<br>REV: TACAGGAGCCTGACACACCATG |
| *Fgf3* | FW: GCAAGCTCTACTGCGCTACCAA<br>REV: CACTTCCACCGCAGTAATCTCC |
| *Ccl2* | FW: GCTACAAGAGGATCACCAGCAG<br>REV: GTCTGGACCCATTCCTTCTTGG |
| *Notum* | FW: CGTGGTACACTCAAGGATGTGC<br>REV: GCCTTATGGCTGTCATGGAAGC |
| *Gh* | FW: TCAGAATGCCCAGGCTGCTTTC<br>REV: ACTGGATGAGCAGCAGCGAGAA |
| *Tsh* | FW: GAGAGTGTGCCTACTGCCTG<br>REV: ACAGACATCCTGAGAGAGTGC |
| *Prl* | FW: CTCAGGCCATCTTGGAGAAG<br>REV: TCGGAGAGAAGTCTGGCAGT |
| *Pomc1* | FW: CCATAGATGTGTGGAGCTGGTG<br>REV: CATCTCCGTTGCCAGGAAACAC |
| *Ccl3* | FW: ACTGCCTGCTGCTTCTCCTACA<br>REV:ATGACACCTGGCTGGGAGCAAA |

Table containing the primer sequences used for qRT-PCR.
